# Software as a Service for the Genomic Prediction of Complex Diseases

**DOI:** 10.1101/763722

**Authors:** Alessandro Bolli, Paolo Di Domenico, Giordano Bottà

## Abstract

In the last decade the scientific community witnessed a large increase in Genome-Wide Association Study sample size, in the availability of large Biobanks and in the improvements of statistical methods to model genomes features. This have paved the way for the development of new prediction medicine tools that use genomic data to estimate disease risk. One of these tools is the Polygenic Risk Score (PRS), a metric that estimates the genetic risk of an individual to develop a disease, based on a combination of a large number of genetic variants.

Using the largest prospective genotyped cohort available to date, the UK Biobank, we built a new PRS for Coronary Artery Disease (CAD) and assessed its predictive performances along with two additional PRS for Breast Cancer (BC), and Prostate Cancer (PC). When compared with previously published PRS, the newly developed PRS for CAD displayed higher AUC and positive predictive value. PRSs were able to stratify disease risks from 1.34% to 25.7% (CAD in men), from 0.26% to 8.62% (CAD in women), from 1.6% to 24.6% (BC), and from 1.4% to 24.3% (PC) in the lowest and highest percentiles, respectively. Additionally, the three PRSs were able to identify the 5% of the UK Biobank population with a relative risk for the diseases at least 3 times higher than the average.

Family history is a well recognised risk factor of CAD, BC, and PC and it is currently used to identify individuals at high risk of developing the diseases. We show that individuals with family history can have completely different disease risks based on PRS stratification: from 2.1% to 33% (CAD in men), from 0.56% to 10% (CAD in women), from 2.3% to 35.8% (BC), and from 1.0% to 34.0% (PC) in the lowest and highest percentiles, respectively. Additionally, the PRSs demonstrated higher predictive performance (AUCs (including age) CAD: 0.81, PC: 0.80, and BC: 0.68) than family history (AUCs (including age) CAD: 0.79, PC: 0.73, and BC: 0.61) in predicting the onset of diseases.

Hyperlipidemia is well known to be associated with higher CAD risk, but a predictive performance comparison between each lipoprotein and CAD PRS has never been assessed. PRS shows higher discrimination capacity and Odds ratio per Standard deviation than LDL, HDL, total cholesterol-HDL ratio, ApoA, ApoB, ApoB-ApoA ratio, and Lipoprotein(a). Comparing the empirical risk distribution between PRS and each lipoprotein, we show that lipoprotein thresholds, currently used in clinical practice, identify a population equal to or smaller than what can be identified with the PRS at the same CAD risk threshold. Moreover, there is not correlation (max *ρ*: 0.137) between PRS and each lipoprotein, indicating that PRS captures different component of CAD etiology and identifies different people at high risk than those identified by lipoproteins, demonstrating to be an invaluable tool in CAD prevention.

One of the major impairment of the PRS usage in clinical practice is the computational complexity needed to calculate per-individual PRSs. Deep bioinformatics expertise is required to run the entire pipeline, from imputing genomic data, through quality control to result visualisation. For these reasons we developed a Software as a Service (SaaS) for genomic risk prediction of complex diseases. The SaaS is fully automated, GDPR complaint and has been certified as a CE marked medical device. We made the SaaS freely available for research purposes. Researchers willing to use the SaaS can contact

## 1 INTRODUCTION

### 1.1 Data is transforming healthcare

An individual’s risk of developing disease results from a complex interaction between their genes and environment. Nevertheless, the collection and analysis of large amounts of data is allowing us to understand these relationships in more detail than ever before and are aiding the development of data-driven approaches to risk prediction that will transform the way that healthcare is provided [1]. From the electronisation of health records to the introduction of wearable devices, developments in digitisation are allowing the collection of biomedical big data at an unprecedented scale ([2] and references therein). At the same time, methods involving advanced analytics like machine learning are leading to a greater understanding of these data [3–7]. The integration of these concepts into healthcare systems are paving the way for a new era of precision medicine. One of the main goal of precision medicine is to use data to identify those at highest risk of developing disease so that interventions and treatments can be targeted at the groups that need them most. This will ensure that finite healthcare resources are used as efficiently as possible and that disease is caught early enough to improve patient outcomes.

Precision medicine is made increasingly possible because we now have the analytical methods and computational power necessary to understand big, complex datasets. These datasets contain clinical outcomes on a large number of people for a variety of diseases and include additional information on physiological, lifestyle and environmental factors, age, sex and family history of disease [8]. When these data are combined and analysed, sophisticated statistical models can find patterns in the combinations of risk factors that lead to disease and provide an estimate of an individual’s risk relative to others in their population with similar values across these factors.

These datasets also increasingly contain genetic or genomic information on individuals as DNA also has a role to play in understanding disease risk. Progress in genomics means that this novel and potentially powerful type of data can be added into disease risk prediction models. The use of genomic data in risk prediction is now possible thanks to the confluence of three major scientific advances:

1. Bigger sample sizes in genome-wide association studies (GWAS) which lead to greater statistical power to identify genetic variants involved with disease [9].
2. Better statistical methods to identify the most predictive set of variants to estimate risk for a given disease [10, 11]
3. The availability of genome-wide data from hundreds of thousands of people linked with thousands of environmental and physiological measurement, such as the UK Biobank, an innovative project developed by the UK Government to collect genetic, health and lifestyle data from 500,000 people that has been made available to scientists and companies. [8]. This magnificent data resource allowed the validation of the algorithms predictive power at an unprecedented scale.

### 1.2 GWAS have highlighted the polygenic architecture of complex diseases

GWAS have been instrumental in understanding the relationship between genetic variation and many complex diseases [9]. The general procedure of a GWAS is to compare the genomes of people with and without a disease (e.g. prostate cancer) or on along a continuous spectrum of trait values (e.g. height). These studies have led to many robust associations of specific genetic variants to a multitude of traits, diseases and disorders. Although this approach does not necessarily find the causative genetic variants contributing to diseases and traits, they provide important clues into their underlying biological basis. The emerging pattern from these studies is that the genetic basis of disease is often, although not exclusively, polygenic [12]. That is, diseases are characterised by contributions from a large number of genetic variants, each of which has an effect of varying magnitude that we can estimate from the GWAS.

Most GWAS have focused on identifying the relationship of a particular type of genetic variant called Single Nucleotide Polymorphisms (SNPs) and disease. SNPs are locations in the human genome where the DNA code is known to vary amongst individuals. The presence of two or more different nucleotides, or alleles, in the DNA sequence at these positions mean that the contribution of an allele to the overall likelihood of disease, or level of a trait, can be estimated. Therefore a GWAS provides two statistics for the alleles at SNP. The first is an estimate of the effect size which is the likelihood to find an allele in the cases rather than the controls. The second is an estimate of the statistical significance of this effect size, which provides a measurement of the confidence we have in the allele’s contribution to the disease. As a rule of thumb, only those SNPs with alleles that gain a genome wide significance *P* value of greater than 5×10*^−^*^8^ are considered as contributing to disease risk (although the procedures described below assess this assumption). The main goal of a GWAS is thus to identify the genetic loci implicated in the aetiology of complex diseases that can subsequently be used to comprehend the molecular mechanisms underlying them.

### 1.3 Large GWAS sample sizes have led to the identification of more genetic variants associated with diseases

Dramatic increases in the number of individuals included in GWAS in recent years are providing the necessary statistical power to uncover statistical significant effect sizes for thousands of genetic variants [13–17]. Because many traits and diseases are polygenic, causal SNPs will each have a small effect size, which are diffcult to estimate accurately with small sample sizes. Although the concepts behind Polygenic Risk Scores (PRS) are not new, with the development of GWASs with sample sizes in the hundreds of thousands, the utility of PRS for predicting disease are being assessed at an unprecedented scale. PRS are computed for an individual by summing the effect alleles weighted by the corresponding values of each effect size to generate a single overall estimate of the genetic risk. Methods for generating PRS and combining them with traditional non-genetic risk factors are developing at pace. However, before the potential of PRS can be realised it is necessary to assess their utility and to understand the challenges of implementing them at scale and the limitations to their use.

### 1.4 Developing a robust procedure for estimating PRS

In its essence, computing an individual’s PRS for a given disease involves multiplying the number of risk alleles a person carries by the effect size of each variant, and then summing these across all risk loci [18]. The accuracy of PRS depends on several conditions [19]. The first is that the GWAS providing the summary statistics for the score – known as the discovery or training dataset – should involve an independent set of samples to those on which the scores are being calculated. Secondly, the amount of variation in disease or trait liability that can be accounted for by the genetic variants used in the PRS, known as SNP heritability, will influence how predictive a PRS will be, which will also be affected by the genetic architecture of the disease [20]. Finally, the sample size of the discovery GWAS will affect how well effect sizes are estimated and therefore affect its accuracy [20]. The best performing PRS will use summary statistics from a discovery GWAS involving hundreds of thousands of independent individuals on a trait with high SNP heritability.

Identifying which SNPs have the best predictive power is a central challenge to developing a robust PRS. There are two main objectives to this effort. The first is to understand at what threshold of statistical significance SNPs should be removed from the score generation algorithm. Because there are correlations between the effect sizes of variants that are close to each other in the genome, the second objective is to explore how to combine evidence across multiple variants.

Procedures have been developed to select subsets of SNPs that rely on looking only at GWAS summary statistics [21]. The simplest approach, known as clumping and thresholding (C+T), iterates between two methodological steps [22]. First, genetic variants are filtered, or clumped, so that only the variants with the highest effect size and that are not in linkage disequilibrium (*LD*) are used. In the second thresholding step, genetic variants with a *P* value larger than a chosen threshold are removed. This process is repeated for different *LD* windows and *P* value thresholds.

A more sophisticated Bayesian approach involves modelling *LD* to shrinks each variant effect size to an extent that is proportional to the *LD* between SNPs [10]. This approach, implemented in the software *LDPred* requires the definition of a tuning parameter *ρ*, which is a statement on a researcher’s prior belief on the proportion of genetic variants assumed to be causal.

Recently, a third method involves machine learning combining C+T and the LASSO statistical procedure, called stacked clumping and thresholding (SCT) has been developed [23]. In SCT, clumping and thresholding are systematically repeated over a four dimensional grid of parameters (comprising LD squared correlation and p-value thresholds). The algorithm generates over 100,000 alternative C+T variants and combines them through a LASSO-based penalized logistic regression.

### 1.5 Validating and testing PRS is possible with the availability of Biobanks

The algorithms outlined above require a validation phase where different PRSs generated with alternative parameter values are validated against an external dataset (Validation dataset). The output of the validation phase is the selection of an optimal PRS, displaying highest predictive performances. The testing phase involves computing the optimal PRS in a test population (Test dataset) and assessing its predictive power in order to confirm its predictive performance and to rule out any possibility of over-fitting that may have occurred during the validation step.

The recent development of PRS has been accelerated by the UK Government, who have made the UK Biobank (UKB) dataset available to researchers and companies [8]. This is a large prospective cohort study that enrolled around 500,000 individuals from across the UK, ranging in age from 40 to 69 years at the time of recruitment and whose genomes have been genotyped and imputed to more than 90 million variants. The astonishing size of the UKB genomic data allowed the building of independent large validation and testing datasets.

### 1.6 Clinical utility and implications of PRS use in the European population

Several studies have assessed the ability of PRS to identify individuals at high risk of developing polygenic diseases. For example, Inouye and colleagues [24] showed that men in the top 20% of PRS distribution reached a threshold of 10% cumulative coronary artery disease (CAD) risk by 61 years of age, ten years earlier than men in the bottom 20% distribution. Additionally it has been shown that for CAD PRS has the higher predictive performance than any of the traditional risk factors (e.g. high cholesterol, family history) used by physicians to decide primary prevention strategies. In a second study, PRS-based models identified 8.0, 6.1, 3.5, 3.2, and 1.5% of the European population at greater than threefold increased risk for Coronary Artery Disease (CAD), Atrial Fibrillation (AT), Type 2 Diabetes (T2D), Inflammatory Bowel Syndrome (IBD), and Breast Cancer (BC), respectively. Most notably for CAD, the prevalence of “been carrier” of high PRS was shown to be 20-fold higher than the prevalence of carriers of the familial hypercholesterolemia mutations that confers the same risk.

### 1.7 Current technological limitations in using PRS

Generating PRS is computationally intensive and so their potential to be used as a tool for precision medicine is currently undervalued. The computers required to generate PRS need hundreds of Gigabytes of RAM and complex computational infrastructures which are extremely difficult to implement and maintain. Additionally, deep bioinformatics expertise is required to run the entire pipeline, from generating genomic data, through quality control to result visualisation. For this reason, analytical laboratories are currently excluded from the possibility to use PRS on a routine basis.

One limitation of GWAS is that they are performed with samples genotyped with microarrays that don’t cover the entire genome, but only a small portion of it. Therefore the causal SNP associated with a phenotype is rarely genotyped, instead the association is attributed to the genotyped SNP in LD with the causal one. However, different ethnic groups are characterised by specific LD patterns, so we can expect that for different ancestries the causal SNP could have a different SNP in LD. For this reason, the SNPs used in a PRS are highly dependent on the genetic structure (i.e. ancestry) of the initial populations used in the GWAS. Since the vast majority of available GWAS is based on population of European descendent (79% of all GWAS partecipants), PRS constructed on these GWAS have the highest predictive power on individuals of the same ancestry. This represents a critical limitation to the mass-scale implementation of PRS in precision medicine and increasing the representation of diverse populations has recently become a higher priority for the research community.

### 1.8 A SaaS for genomic risk prediction addresses current limitations in PRS utilisation

We present a Software as a Service (SaaS) for genomic risk prediction based on PRS. A SaaS is a software distribution model in which a third-party provider hosts applications and makes them available to users over the Internet. The SaaS takes as input genome-wide data of an individual in the form of a text file. This text file contains an individual’s genetic information, which can be in one of several different formats depending on the microarray and Next-generation sequencing platform used to generate the data. Both high coverage and low pass sequencing are currently available on the market and can be easily used as input files. Once files are uploaded into the SaaS, they are loaded to a secure network file system hosted in the cloud where data are processed. Data processing involves data conversion to a common format, imputation of uncalled variants, quality control and finally PRS calculation and visualisation.

## 2 CASE STUDIES

### 2.1 Overview

In this section we describe the development and validation of Polygenic risk scores (PRSs) for three diseases: coronary artery disease (CAD), breast cancer (BC) and prostate cancer (PC). We also assessed how a published PRS for CAD based on millions of SNPs [21] performed with different subsets of SNPs. We found that although a large fraction of SNPs had an effect size that is close to zero, including them in the PRS maximized the prediction performance of the score.

For all diseases, we show that PRS can identify a large proportion of the population that is at least three times more likely of developing the disease compared to the average. We also show that PRSs for CAD, PC, and BC maintain their high stratification power, even in individuals already considered at higher risk of disease due to having at least one first-degree relative with a family history of CAD, PC or BC. Moreover, we found that PRSs are better than family history in terms of prediction performances for each of the three diseases.

Finally for CAD we compared the predictive performance of PRS with those of the lipoproteins currently used in clinical practise. We found that PRS is a stronger risk factor than lipoproteins and is also orthogonal identifying different individuals than those identified by lipoproteins above clinically relevant thresholds.

### 2.2 Scientific Methodology

We used GWAS summary statistics generated from *training* datasets. For each disease considered, we built a PRS using algorithms that identify the optimal subset of SNPs and their effect sizes from the GWAS summary statistics to maximize the predictive performances in a *validation* dataset. Next, we tested whether the PRSs with the highest predictive power in the validation dataset had the same performance in an independent *testing* dataset. Finally, we compared the predictive power of newly developed PRS with previously published PRS using the *testing* dataset.

#### 2.2.1 Discovery and validation of PRS for CAD, PC, and BC

To build the PRSs, we used the SCT method of Privè and colleagues [23], implemented in *R* [25]. The method uses summary statistics from published GWAS, genetic and clinical data from the validation dataset. The interim release of UK Biobank (i.e. individuals genotyped through batches from 1 to 22) was used as a validation dataset. The UK Biobank fields used to define cases of CAD, BC, and PC are reported in Table 1. SCT uses per-SNP effect sizes and *P* values to perform repeated clumping and thresholding (C+T) over a four dimensional grid of parameters (comprising *LD* squared correlation, *P* value threshold, clumping window size, and imputation quality).

**Table 1:** Lists of UK Biobank fields and codes defining cases of CAD, BC, and PC.

| Coronary Artery Disease (CAD) |  |  |
| --- | --- | --- |
| UK Biobank Field | Description | Codes |
| 20004 | Self Reported Operation code | 1070, 1095, 1523 |
| 20002 | Self Self-reported non-cancer illness | 1075 |
| 41202 | Diagnoses - main ICD10 | I21X, I22X, I23X,<br>I241, I252 |
| 41204 | Diagnoses - Secondary ICD10 |  |
| 41200 | Operative procedures - main OPCS4 | K401-K404,<br>K411-K414,<br>K451-K455,<br>K491-K492<br>K498-K499, K502,<br>K751-K754,<br>K758-K759 |
| 41210 | Operative procedures - secondary OPCS4 |  |
| Prostate Cancer (PC) |  |  |
| UK Biobank Field | Description | Codes |
| 20001 | Self-reported cancer code | 1044 |
| 41202 | Diagnoses - main ICD10 | C61, D075 |
| 41204 | Diagnoses - Secondary ICD10 |  |
| 40001 | Underlying cause of death ICD10 |  |
| 40002 | Contributory causes of death ICD10 |  |
| 40006 | Type of cancer ICD10 |  |
| Breast Cancer (BC) |  |  |
| UK Biobank Field | Description | Codes |
| 20001 | Self-reported cancer code | 1002 |
| 41202 | Diagnoses - main ICD10 | C500-C509,<br>D050-D051,<br>D057, D059 |
| 41204 | Diagnoses - Secondary ICD10 |  |
| 40001 | Underlying cause of death ICD10 |  |
| 40002 | Contributory causes of death ICD10 |  |
| 40006 | Type of cancer ICD10 |  |

Overall, SCT generated 123,200 alternative C+T configurations, each of which was used to compute a corresponding PRS in the validation dataset. PRS scores were generated as the sum of the genotype dosage of each risk allele at each SNP weighted by its respective effect size. These PRSs were then used as the predictive variables in a LASSO-penalized logistic regression model with disease phenotype as the binary response variable, generating a regression coefficient for each C+T configuration. The stacking phase followed, where effect sizes and regression coefficients of the 123,200 alternative C+T configurations were linearly combined to generate a final optimal panel of SNPs for the PRS. We define a PRS panel as a 3-column table of SNPs, effect alleles, and corresponding effect sizes. For CAD, PC, and BC, we utilised published GWAS summary statistics from Nikpay et al., [13], Schumacher et al., [26], and Michailidou et al., [15], respectively.

#### 2.2.2 Evaluating PRSs predictive performances

We assessed the predictive performances of CAD, PC and BC PRSs, on an independent testing dataset comprising the second release of UK Biobank (i.e. individuals genotyped in batches from 23 to 95). As a comparison, we also re-calculated the published PRS panels from Khera[22] (PRS for CAD and BC) from Inouye [24] (PRS for CAD), from Mavaddat [27] (for BC), and from Schumacher[14] (for PC) in this testing dataset and compared their predictive performance with the PRSs developed in this study. PRS values were used as predictive variable in a logistic regression model. The logistic regression model comprised additional covariates such as: age, gender, genotyping array, and the first 4 principal components (PCs) of ancestry. The PC and BC analyses were restricted to male and female participants, respectively. We assess the predictive performance of each PRS panel by computing the Area Under the Curve (AUC) of the Receiver Operating Characteristic (ROC) curve and the Positive predictive value (PPV) using as a treshold the top 3% of the PRS distribution.

We compared results to previously published PRS for CAD [22, 24], BC [22, 28], and PC [14] downloaded from the Broad Institute Cadiovascular Disease knowledge portal^*^or from the supplementary information of related papers^†^

### 2.3 Coronary Artery Disease

CAD is a disease caused by the narrowing or blockage of the coronary arteries and is usually caused by atherosclerosis, a hardening of the arteries. Whilst environmental and lifestyle factors can modulate an individual’s risk of developing CAD [29], a genetic component is also known [13], making a perfect candidate for the investigation of PRS predictive performance.

#### 2.3.1 Variants with small effect sizes play a role in Genome-Wide PRS

Two PRS for CAD have recently been published. Khera and colleagues [22] used the *LDpred* algorithm [10] to develop a CAD PRS using 6.6 million SNPs, while Inouye and colleagues [24] aggregated three PRS for CAD to generate a *metaPRS* using 1.7 million SNPs. Using *LDpred*, Khera and colleagues obtained a PRS with the best predictive performance with a parameter value that indicates that 0.1% of the variants in the analysis are causal. This implies that only the 0.1% of the 6.6 million SNPs in the PRS should have an effect on the prediction, while the remaining 99.9% have an effect size close to zero (of about 3×10*^−^*^6^).

Given that in this analysis the vast majority of SNPs were estimated to have minimal or no effect on polygenic risk, doubt has been cast on the utility of including this large fraction of low-weight SNPs in PRSs [30]. We addressed the effect of including different numbers of SNPs in Khera PRS comparing predictive power across progressively smaller subsets of SNPs (Table 2): the full PRS (6.6 million SNPs), a PRS made by the top 1% of SNPs with highest effect sizes (66,300 SNPs), a second PRS constituted by the top 0.1% of SNPs with highest effect sizes (6630 SNPs), as well as a PRS generated with genome-wide significant SNPs only (*P* value < 5×10*^−^*^8^, corresponding to 74 SNPs). AUC [31] and Positive Predictive Value (PPV) at 3% were calculated for each PRS in the testing dataset. Table 2 shows that a decrease in the number of SNPs used in a PRS is matched by a corresponding decrease in its discriminatory ability in both AUC and PPV. This finding demonstrates that even a set of low weight SNPs can play a crucial role in a more accurate risk prediction.

**Table 2:** SNP subsets of PRS for CAD from Khera et al assessed in this study. Khera full refers to the whole PRS for CAD developed by Khera et al^22^. Khera 1% refers to the PRS generated with the 1% of genetic variants with highest effect sizes from Khera et al. Khera 0.1% refers to the PRS generated with the 0.1% of genetic variants with highest effect sizes from Khera et al. Khera 74 refers to the PRS generated with genome-wide significant SNPs only as described in Khera et al. For each PRS, Table 2 shows the number of genetic variants composing the PRS (**SNPs in PRS**), the predictive performances quantified as AUC values and 95% confidence intervals (**AUC (95% CI)**), the positive predictive values in the top 3% of PRS distributions (**PPV (3%)**), and the number of CAD cases in the top 3% of PRS distributions (**Cases in top 3%**).

| PRS panel | SNPs in PRS | AUC (95% CI) | PPV (3%) | Cases in top 3% |
| --- | --- | --- | --- | --- |
| Khera full | 6630150 | 0.805 (0.798–0.812) | 12.35 | 1031 |
| Khera 1% | 66300 | 0.798 (0.792–0.805) | 11.31 | 945 |
| Khera 0.1% | 6630 | 0.794 (0.788–0.801) | 10.88 | 909 |
| Khera 74 | 74 | 0.789 (0.784–0.797) | 9.63 | 804 |

#### 2.3.2 Development of a new CAD PRS

We next assessed the predictive performance of our new PRS in the testing dataset (population sample size of 278000 individuals). The SCT PRS displayes higher predictive performances (AUC: 0.808, PPV at 3%: 13.06%) than the PRSs from Khera et al[22] (AUC: 0.805, PPV at 3%: 12.35%) and Inouye et al[24] (AUC: 0.805, PPV at 3%: 12.5%) (see Table 3). Notably, the final CAD PRS developed using SCT was composed of only 300,000 genetic variants, a number that corresponds to only the 5% and 17% of the SNPs of CAD PRS from Khera and Inouye, respectively.

**Table 3:** List of the PRSs for CAD assessed in this study. Khera full refers to the whole PRS for CAD developed by Khera et al. Inouye full refers to the whole PRS for cad developed by Inouye et al. SCT refers to the PRS for CAD developed in this paper with the SCT algorithm. SCT + Khera refers to the PRSs generated by combining SCT and full Khera PRS ^(1)^. SCT + Inouye refers to the PRSs generated by combining SCT and full Inouye PRS^(1)^. For each PRS, Table 3 shows the number of genetic variants composing the PRS (**SNPs in PRS**), the predictive performances quantified as AUC values and 95% confidence intervals (**AUC (95% CI)**), the positive predictive values in the top 3% of PRS distributions (**PPV (3%)**), and the number of CAD cases in the top 3% of PRS distributions (**Cases in top 3%**). ^(1)^For SNPs with effect sizes from both SCT and the second panel (khera or Inouye), effect sizes from SCT were taken.

| PRS panel | SNPs in PRS | AUC (95% CI) | PPV (3%) | Cases in top 3% |
| --- | --- | --- | --- | --- |
| Khera full | 6630150 | 0.805 (0.798–0.812) | 12.35 | 1031 |
| Inouye full | 1745180 | 0.805 (0.799–0.812) | 12.5 | 1044 |
| SCT | 291969 | 0.808 (0.8–0.815) | 13.06 | 1091 |
| SCT+ khera <sup>(1)</sup> | 6699370 | 0.808 (0.801–0.814) | 12.99 | 1085 |
| SCT+ Inouye <sup>(1)</sup> | 1920136 | 0.810 (0.803–0.816) | 13.36 | 1116 |

In light of our finding outlined above, that a larger number of SNPs - even if with low effect sizes - improve the predictive performance of a PRS, we asked whether integrating the large SNP sets from the Khera or Inouye studies to our CAD PRS could further improve its predictive performance. We found that the addition of SNPs from Inouye to the SCT PRS led to a new CAD PRS (denoted as SCT + Inouye or SCT-I) with further improved predictive performance, as quantified by an increased value of the AUC (0.81) and the PPV at 3% (13.36%) (Table 3). This finding highlights the highly polygenic nature of this common disease and is also theoretically consistent with the omnigenic model proposed by Pritchard and colleagues [32]. The omnigenic models describes how genes expressed in disease-relevant cells are able to affect core genes directly involved in the disease though a complex regulatory networks. To combine the SNPs in the CAD PRS, three strategies were used:

1. Inverse variance-weighted average method (IVW), where SNPs common in two PRS panels are aggregated by performing the weighted sum of each SNP effect size, weighted by the inverse of each PRS panel’s variance [29].
2. Following the approach of Inouye et al. [24], where SNP effect sizes from two PRS panels are summed after being divided by the standard deviation of each PRS panel.
3. Aggregating SNPs effect sizes from different PRS panels without any normalization (reported in the Table above).

The meta-PRS of SCT+Khera and SCT+Inouye generated through the first two methodologies didn’t provide any further increase in the predictive performances, respect to the original PRS panel. This was due to the fact that the standard deviation of the effect sizes of the SCT panel is several times higher than those of Khera and Inouye. For this reason, normalization methods 1 and 2 tended to under-estimate effect sizes from SCT in favor of those of Khera or Inouye. This un-balance did not occur when we aggregated effect sizes without normalization. In this latter case, we observed an increase in the AUC of the SCT+Inouye PRS respect to the two original PRSs. In Table 3, the metrics of the PRSs derived from the un-normalized method are shown.

#### 2.3.3 Predictive performance of the newly developed CAD PRS: SCT-I

The following analyses are based on the SCT-I CAD PRS that showed the highest predictive performance (Table 3). We computed the PRS of the individuals in the testing dataset and plotted the distributions of the scores for CAD cases and controls (Figure 1A). The distributions are both gaussian, with CAD cases showing a greater median PRS than controls (median: 0.52 and −0.03, respectively) and an AUC of 0.81.

**Figure 1:**
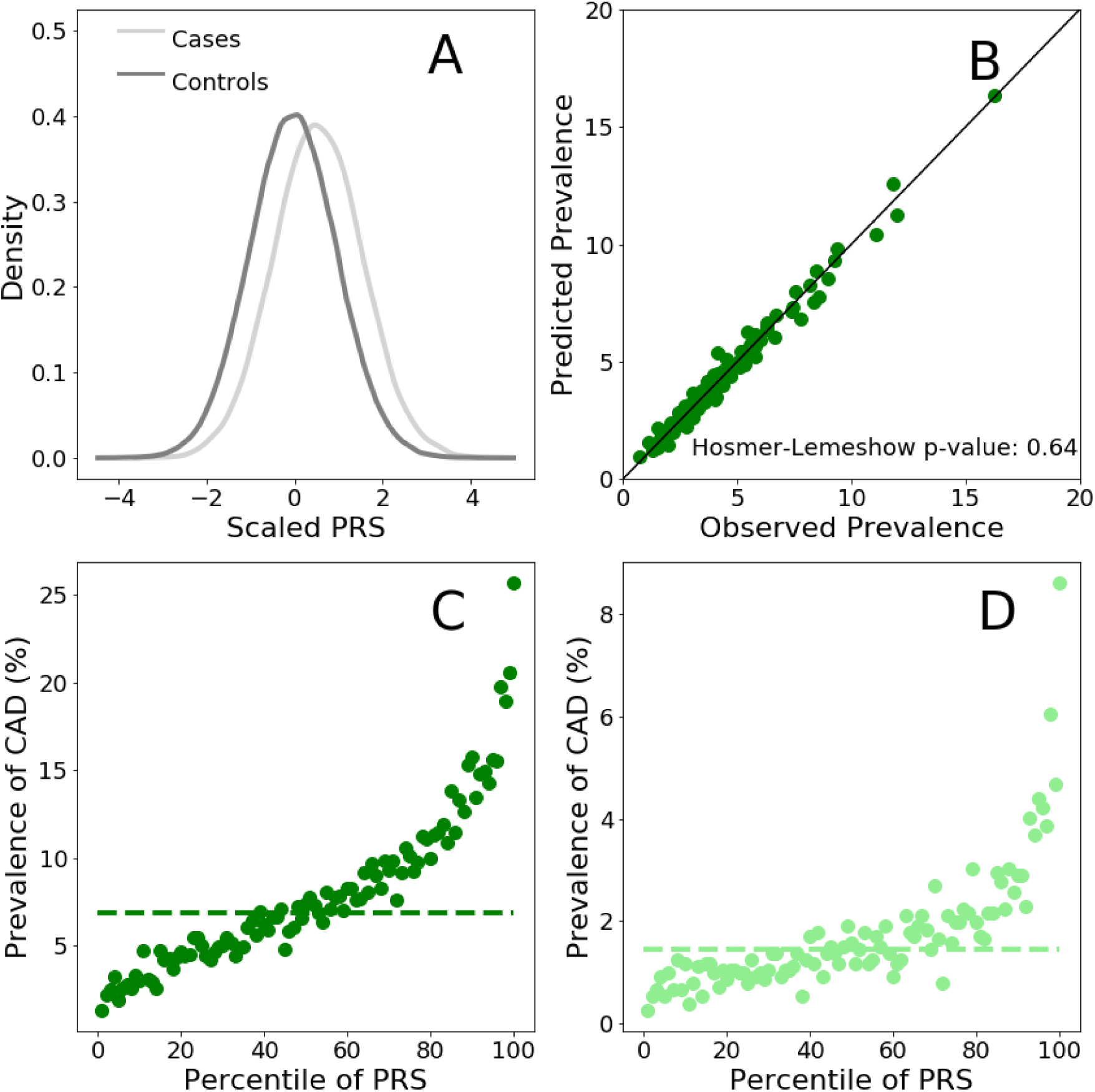
Risk for CAD according to the SCT-I PRS panel. Panel A. Distributions (scaled to a mean of 0 and to a standard deviation of 1) of the PRS score for CAD cases and controls in the testing population. **Panel B.** Comparison between the observed and predicted CAD prevalences. Observed prevalence has been calculated as the per-percentile prevalence of CAD in the PRS distribution. Predicted CAD prevalence was calculated for each individual using a logistic regression model with PRS as predictive variables. Within each percentile of the PRS distribution, CAD probability was averaged and this returned the predicted prevalence of CAD. **Panel C.** Prevalence of CAD per percentile of the PRS distribution calculated in the men testing population. **Panel D.** Prevalence of CAD per percentile of the PRS distribution calculated in the women testing population. Dashed horizontal lines: CAD prevalence of the average of the PRS distributions (defined as between the 40% and the 60% percentiles) for men (dark green) and women (light green)

We next evaluated the ability of the SCT-I PRS to stratify CAD risk separately for sub-populations of men (about 126000 individuals) and women (about 152000 individuals) in the testing dataset. We divided the two PRSs distributions into percentiles and computed the prevalence of CAD in each percentile. Here we use disease prevalence in the testing dataset as a measure of the risk of developing CAD. Risk stratification for men and women in the testing dataset are shown in Figure 1C and Figure 1D, respectively. CAD risk rises sharply as PRS percentile increases, ranging from 1.34% to 25.67% (for men) and from 0.26% to 8.62% (for women), for the lowest and highest percentiles, respectively.

For each sex we estimated the relative increased risk, which is the ratio between the prevalence at the top 5% of the PRS distribution and the prevalence in the average of the distribution (defined as between the 40% and the 60% percentiles). For men, the relative risk in the top 5% is 3 times higher than the average while for women this value rises to 4. This means that the CAD SCT-I PRS is able to detect individuals with a three fold relative risk of developing CAD which is comparable to that conferred by rare highly penetrant familial hypercholesterolemia mutations [33].

Above, we showed how the SCT-I PRS can stratify the empirical risk of CAD in a testing dataset with known disease prevalence. However, the clinical value of a PRS implies its ability to predict the risk of disease in a well calibrated model. To assess model calibration we compared the predicted prevalence values with observed ones. For each individual within the testing dataset, the probability of having the disease was calculated using a logistic regression model with the PRS score as predictive variable. The predicted prevalence of CAD within each percentile of the PRS distribution was calculated as the average probability in each percentile. For all percentiles, predicted CAD prevalence was plotted against the corresponding values of observed prevalence (Figure 1B). Figure 1B shows that the values of observed and predicted CAD prevalence are in excellent agreement as demonstrated by the localization of the points of the bisector of the graph. We also tested the level of agreement between the predicted and observed prevalence through the Hosmer-Lemeshow (HL) test. This is a goodness of fit testing for logistic regression, especially for risk prediction models. Specifically, the HL testing calculates if the observed prevalence matches the predicted prevalence in population subgroups represented by PRS percentiles. The non-signicant p-value generated by the HL testing (Figure 1B) implies that there is no statistical evidence of a deviation between observed and predicted prevalence values, thus confirming the good fit of the calibration that can be observed in (Figure 1B).

#### 2.3.4 PRS is more effective at predicting CAD risk than family history

Family history of heart disease is a well-recognized risk factor and prospective studies demonstrate a consistent association with the disease [34]. Family history can be easily and systematically queried in the clinical setting. In this section, we considered the relationship between two risk factors for CAD: family history and PRS. In particular we wanted to answer the following questions:

1. Can SCT-I PRS stratify risk in people with family history?
2. Is SCT-I PRS a better predictor than family history?
3. Does prediction performance increase if a combination of family history and PRS is used?

We computed PRS for CAD in cases and controls for those individuals in the testing dataset with at least one first-degree relative with a history of heart disease. PRS distributions for cases and controls are shown in Figure 2A. Both distributions are gaussian with cases having a higher median value than controls (median: 0.60 and 0.08, respectively). This shows that the good discriminatory ability of the SCT-I PRS is maintained even in individuals already considered at CAD risk based on family history.

**Figure 2:**
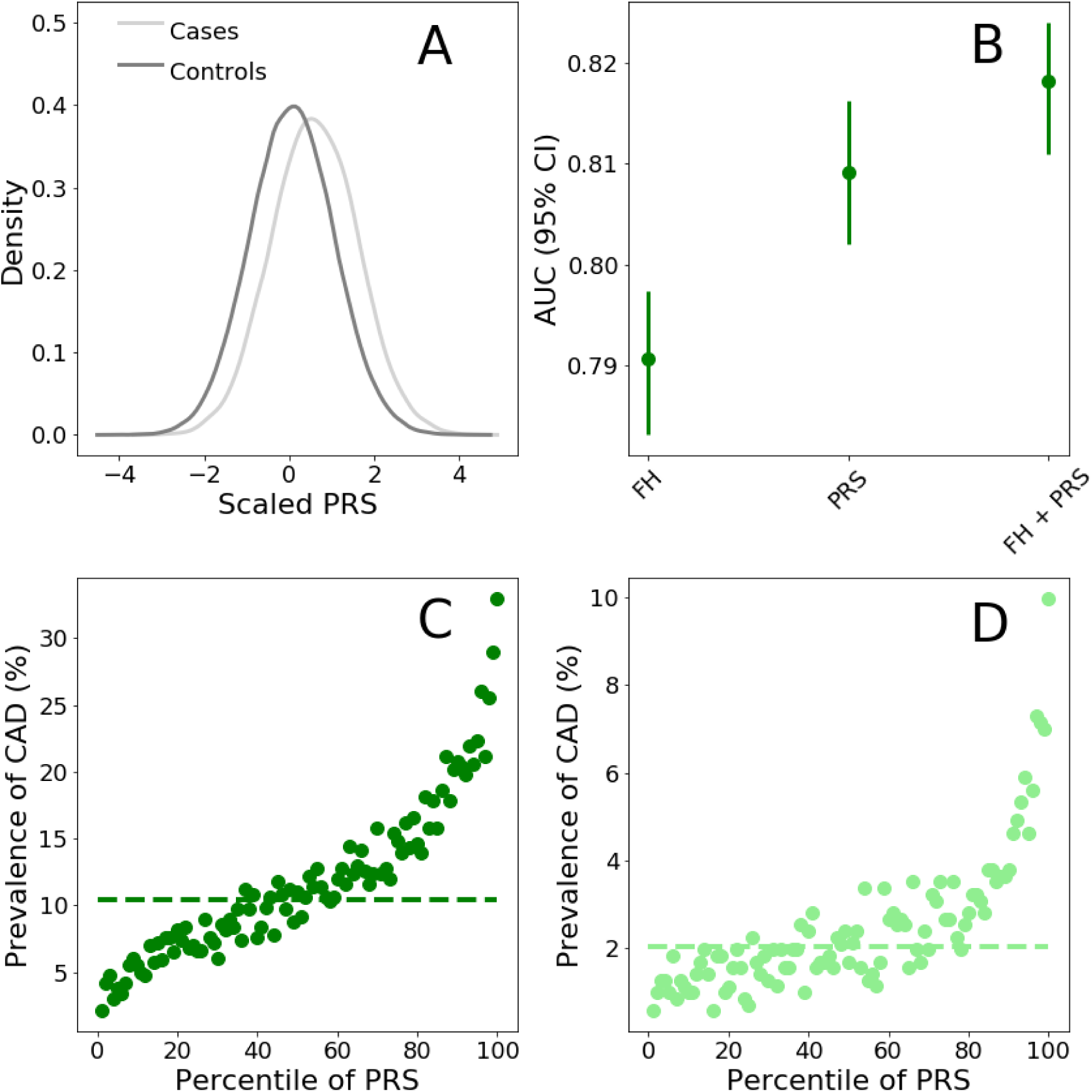
Risk for CAD according to the SCT-I PRS panel in presence of Family history of heart disease. Panel A. Distributions (scaled to a mean of 0 and to a standard deviation of 1) of the PRS score for CAD cases and controls with at least one first-degree relative with a history of heart disease. **Panel B.** AUC values were calculated on the testing dataset with logistic regressions using family history of heart disease (FH), PRS calculated with the SCT-I PRS (PRS), or both (FH + PRS) as explanatory variables. The response variable of the logistic regression model was absence/presence of CAD. The logistic regression model comprised additional covariates such as: age, gender, genotyping array, and the first 4 principal components (PCs) of ancestry. **Panel C.** Prevalence of CAD per percentile of the PRS score distribution calculated in the testing dataset for men with at least one first-degree relative with history of heart disease. **Panel D.** Prevalence of CAD per percentile of the PRS distribution calculated in the testing population for women with at least one first-degree relative with history of heart disease. Dashed horizontal lines: CAD prevalence of the average of the PRS distributions (defined as between the 40% and the 60% percentiles) for men (dark green) and women (light green)

We then evaluated the ability of the SCT-I PRS to stratify CAD risk in the sub-populations of men and women with at least one first-degree relative with a history of heart disease. CAD risk stratification for men and women with family history of heart disease are shown in Figure 2C and 2D, respectively. Even with individuals considered at higher risk based on family history, the SCT-I PRS was able to further stratify CAD risk over a range of values comprised between 2.10% and 33% (for men) and between 0.56% and 10% (for women), for the lowest and highest percentiles, respectively. For both men and women, average prevalence was higher in individuals with family history than in the general population for any percentile considered. For men with family history, the relative risk in the top 5% is 3 folds higher than the average while for women this value rises to 4.

Lastly, we assessed the discrimination ability of family history, PRS, and the combination of the two, by computing AUC. Figure 2B shows that PRS displays a higher AUC value than family history (AUC PRS: 0.810 (0.803-0.815), AUC family history: 0.791 (0.783-0.797)) and has therefore better capacity to discriminate between CAD cases and controls. When both risk factors are combined, the predictive performances improves further (AUC: 0.817 (0.811-0.824)).

These findings demonstrate that family history and PRS capture different components of the risk of CAD and family history cannot be considered in isolation without further PRS risk stratification.

#### 2.3.5 SCT-I for CAD is more predictive than and orthogonal to risk-enhancing lipoproteins

Low-density lipoprotein cholesterol (LDL-C) is recognized as a primary lipid risk factor of CAD. Additionally, several major guidelines suggest to consider high-density cholesterol (HDL-C), and lipid ratios such total cholesterol/HDL-C (tCHL/HDL-C) as additional risk factors [35, 36]. There are also evidences suggesting apolipoproteins (Apo) as effective risk factors of cardiovascular disease. Indeed, Apolipoprotein B (ApoB), that is a proxy of the number of potentially atherogenic lipoprotein particles, and Apolipoprotein A-I (ApoA), which reflects antiatherogenic HDL, may be better indicators of cardiovascular risk. In particular, the ApoB/ApoA ratio has been shown to be strongly associated with the risk of myocardial infarction and stroke [37–39]. Here, we compare the predictive performance and assess correlation of the SCT-I PRS with several plasma lipoproteins. As plasma risk factors for CAD we took into consideration the following: LDL cholesterol (LDL-CHL) (Figure 3), HDL-cholesterol (HDL) (Figure 4), the total cholesterol HDL ratio (tCHL-HDL) (Figure 5), ApoA (Figure 6), ApoB (Figure 7), the ApoB ApoA ratio (ApoB-ApoA) (Figure 8), as well as Lipoprotein(a) (Lp(a)) (Figure 9).

**Figure 3:**
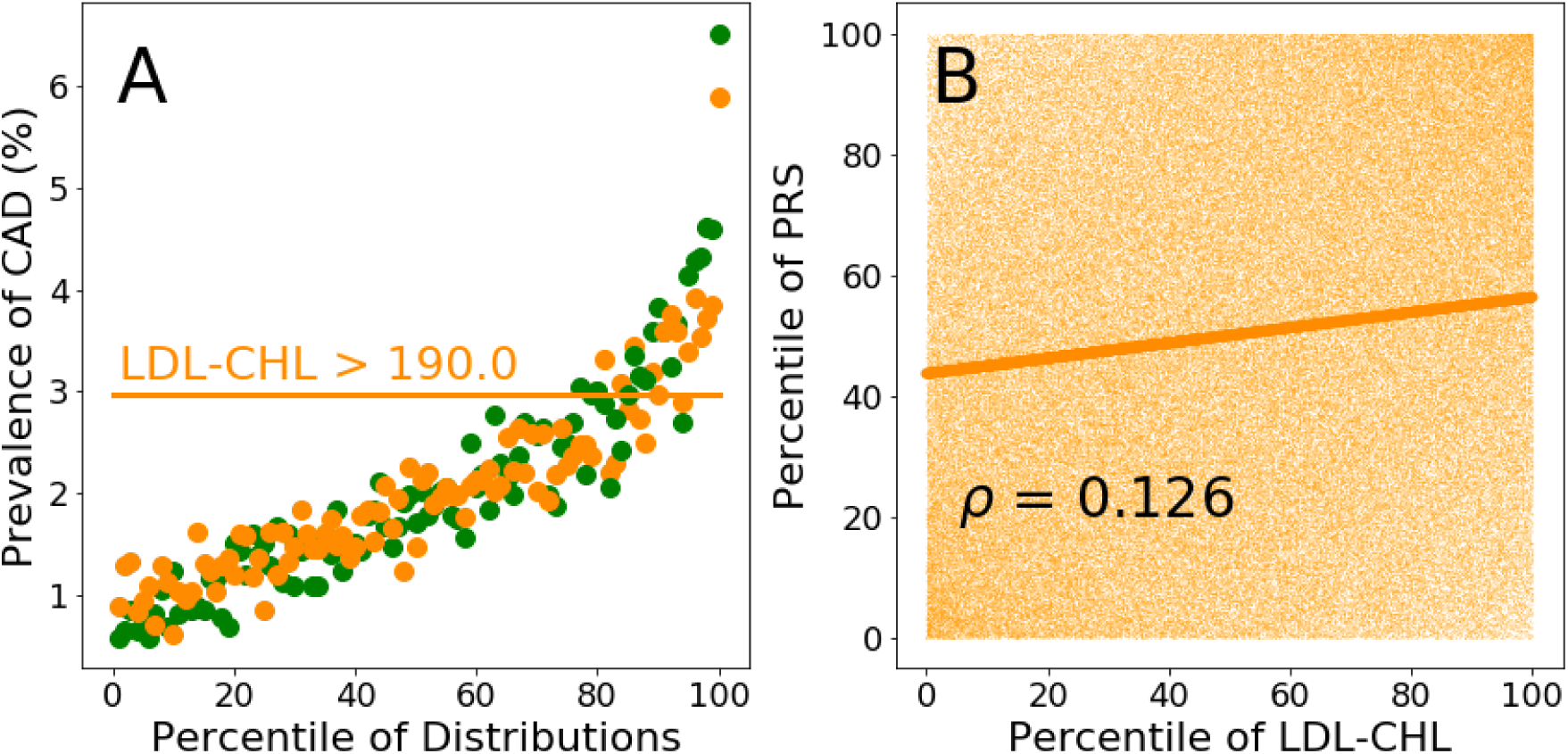
Predictive performance of the SCT-I PRS and LDL-cholesterol. Panel A. Prevalence of CAD per percentile of the PRS (green dots) and LDL-C (orange dots) distributions calculated in the testing population. The horizontal line represents the value of CAD prevalence (3%) at which the LDL-cholesterol distribution reaches a value of 190 (in correspondence of the 90^th^ percentile of its distribution). The same CAD prevalence value is reached by the PRS distribution at its 76^th^ percentile. **Panel B.** Scatter plot of the percentiles of the PRS distribution plotted against the percentiles of the LDL-cholesterol distribution. Orange continuous line: linear regression of the scatter plot

**Figure 4:**
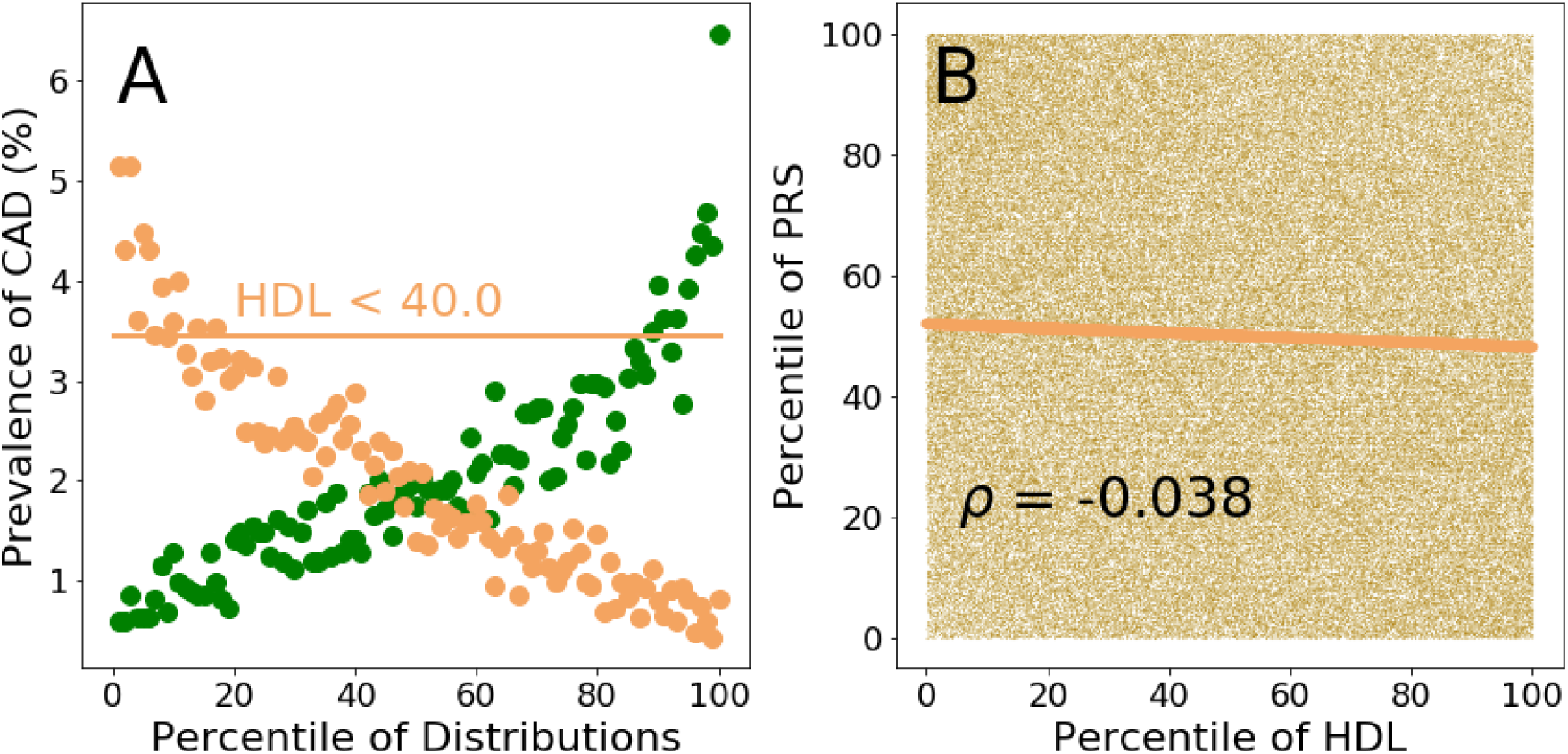
Predictive performance of the SCT-I PRS and HDL cholesterol (HDL). Panel A. Prevalence of CAD per percentile of the PRS (green dots) and HDL (sandy brown dots) distributions calculated in the testing population. The horizontal sandy brown line represents the value of CAD prevalence (3.5%) at which the HDL distribution reaches a value of 40 mg/dl (in correspondence of the 11^th^ percentile of its distribution). The same CAD prevalence value is reached by the PRS distribution at its 88^th^ percentile. **Panel B.** Scatter plot of the percentiles of the PRS distribution plotted against the percentiles of the HDL distribution. Sandy brown continuous line: linear regression of the scatter plot.

**Figure 5:**
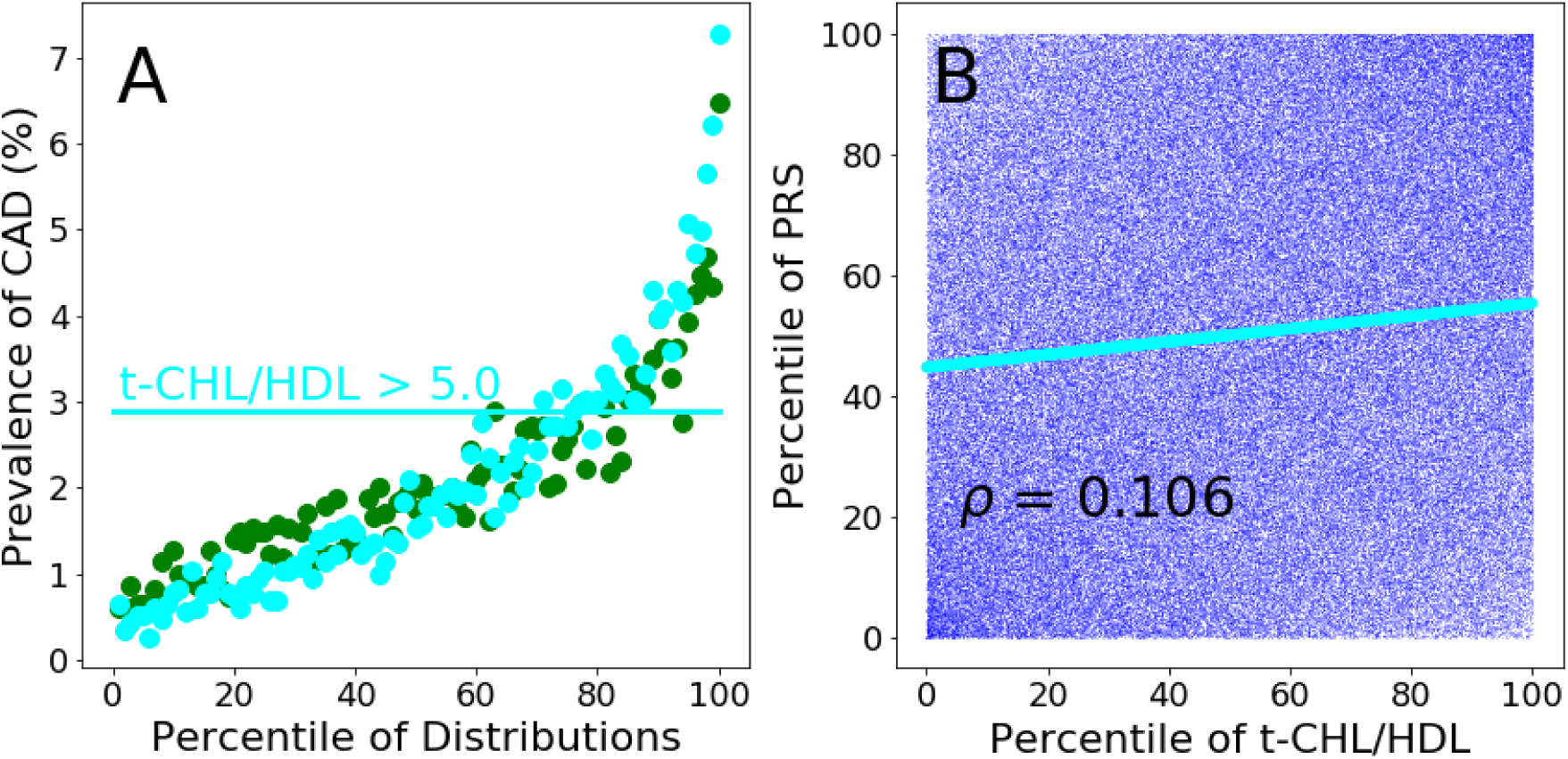
Predictive performance of the SCT-I PRS and total cholesterol-HDL ratio (t-CHL-HDL). Panel A. Prevalence of CAD per percentile of the PRS (green dots) and t-CHL-HDL (cyan dots) distributions calculated in the testing population. The horizontal line represents the value of CAD prevalence (2.9%) at which the t-CHL-HDL distribution reaches a value of 5.0 (in correspondence of the 75^th^ percentile of its distribution). The same CAD prevalence value is reached by the PRS distribution at its 76^th^ percentile. **Panel B.** Scatter plot of the percentiles of the PRS distribution plotted against the percentiles of the t-CHL-HDL distribution. Cyan continuous line: linear regression of the scatter plot.

**Figure 6:**
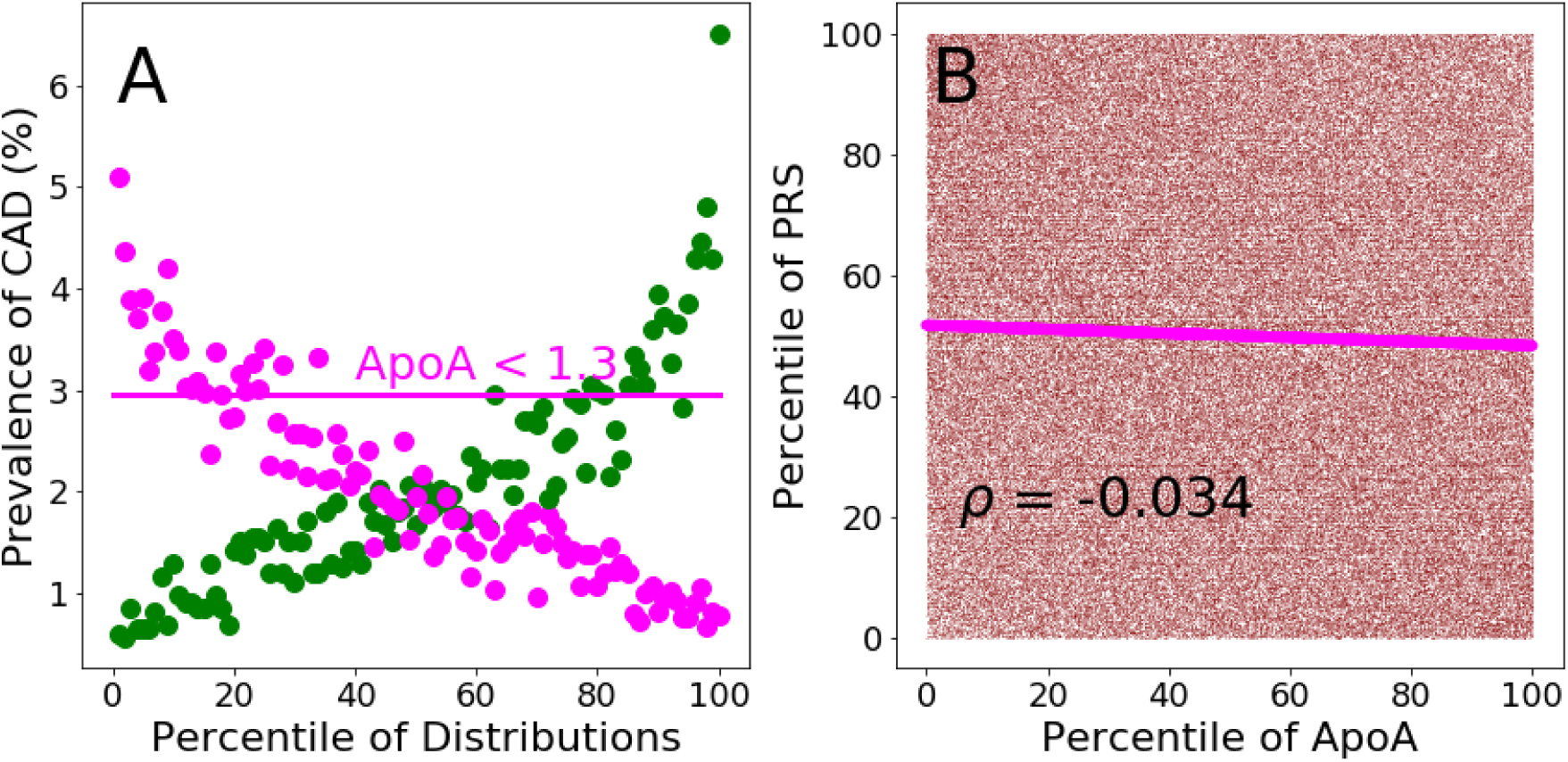
Predictive performance of the SCT-I PRS and Apolipoprotein A (ApoA). Panel A. Prevalence of CAD per percentile of the PRS (green dots) and ApoA (magenta dots) distributions calculated in the testing population. The horizontal line represents the value of CAD prevalence (3%) at which the ApoA distribution reaches a value of 1.3 g/L (in correspondence of the 16^th^ percentile of its distribution). The same CAD prevalence value is reached by the PRS distribution at its 62^th^ percentile. **Panel B.** Scatter plot of the percentiles of the PRS distribution plotted against the percentiles of the ApoA distribution. Magenta continuous line: linear regression of the scatter plot.

**Figure 7:**
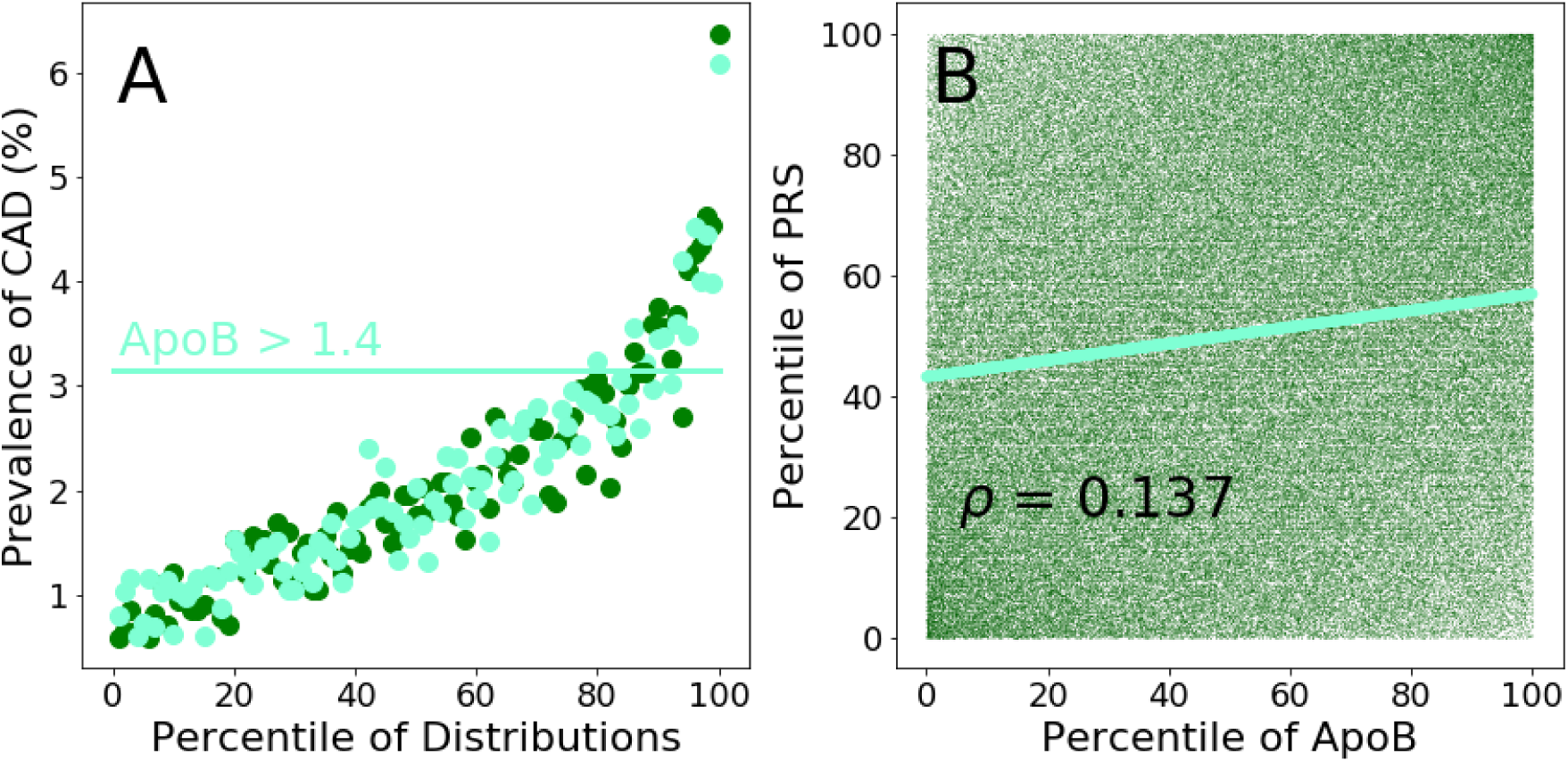
Predictive performance of the SCT-I PRS and Apolipoprotein B (ApoB). Panel A. Prevalence of CAD per percentile of the PRS (green dots) and ApoB (aquamarine dots) distributions calculated in the testing population. The horizontal line represents the value of CAD prevalence (3.1%) at which the ApoB distribution reaches a value of 1.4 g/L (in correspondence of the 89^th^ percentile of its distribution). The same CAD prevalence value is reached by the PRS distribution at its 85^th^ percentile. **Panel B.** Scatter plot of the percentiles of the PRS distribution plotted against the percentiles of the ApoB distribution. Aquamarine continuous line: linear regression of the scatter plot.

**Figure 8:**
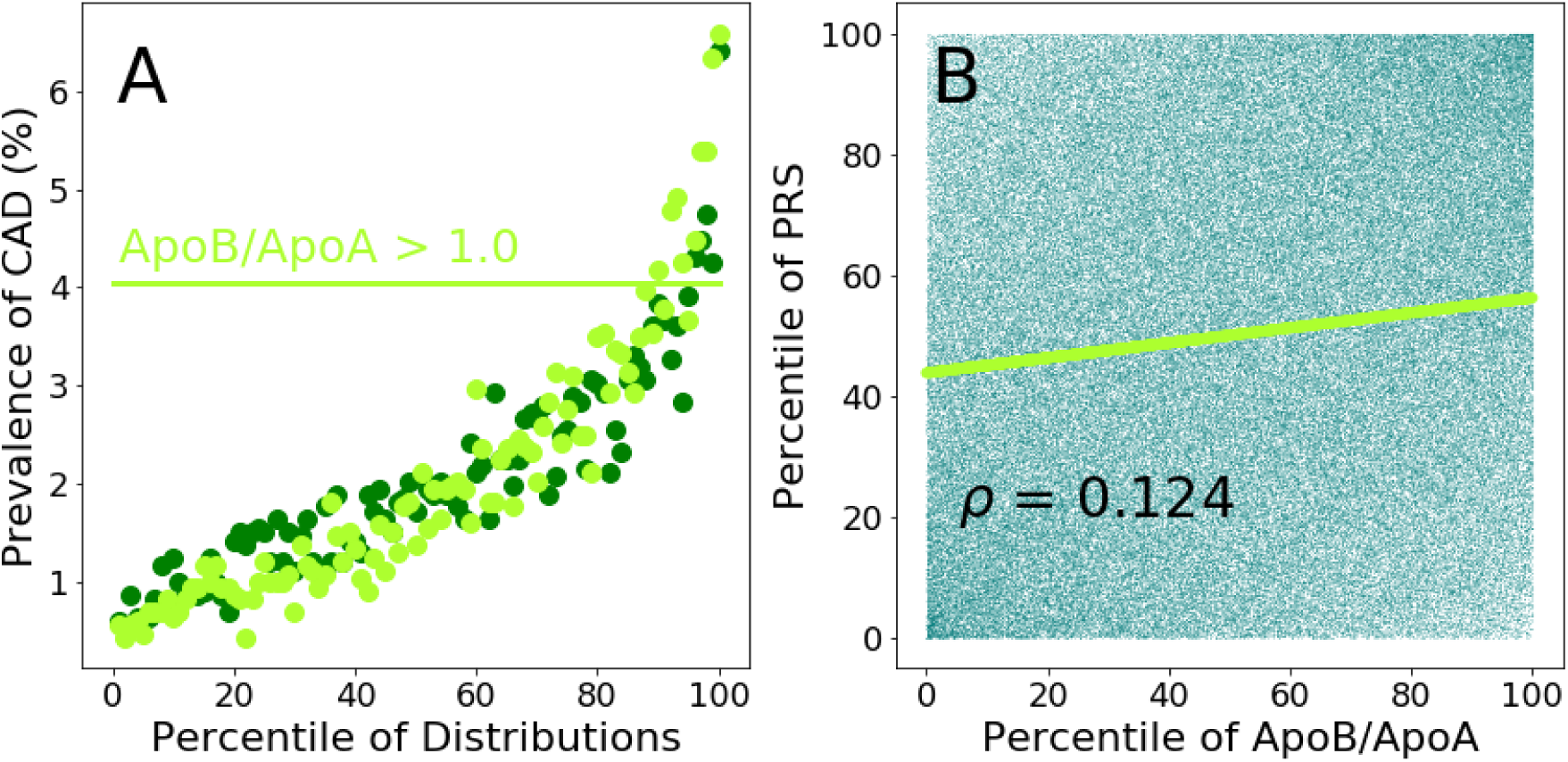
Predictive performance of the SCT-I PRS and Apolipoprotein B-Apolipoprotein A ratio (ApoB-ApoA). Panel A. Prevalence of CAD per percentile of the PRS (green dots) and ApoB-ApoA (greenyellow dots) distributions calculated in the testing population. The horizontal line represents the value of CAD prevalence (4.0%) at which the ApoB-ApoA distribution reaches a value of 1.0 (in correspondence of the 90^th^ percentile of its distribution). The same CAD prevalence value is reached by the PRS distribution at its 95^th^ percentile. **Panel B.** Scatter plot of the percentiles of the PRS distribution plotted against the percentiles of the ApoB-ApoA distribution. Greenyellow continuous line: linear regression of the scatter plot.

**Figure 9:**
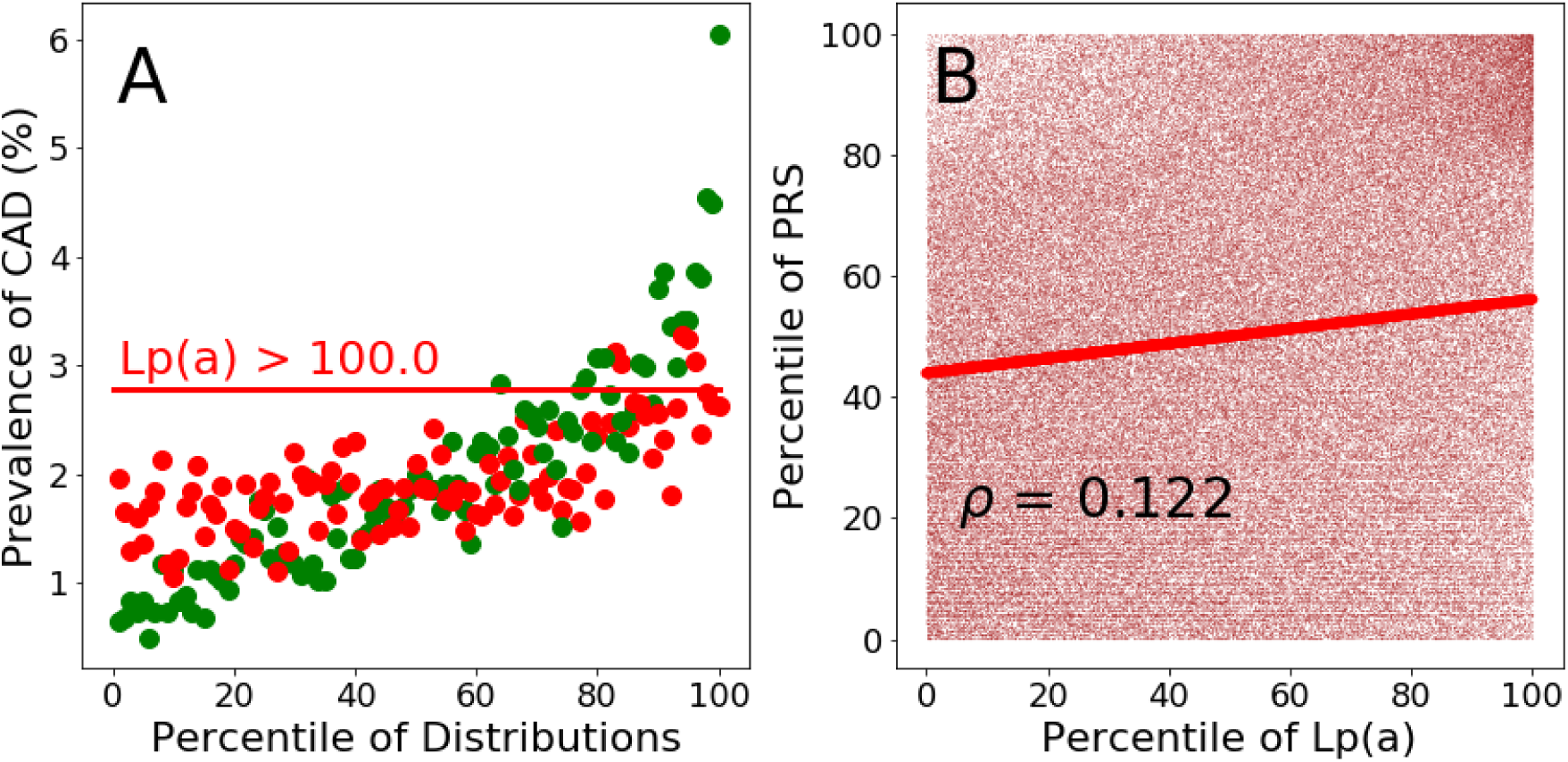
Predictive performance of the SCT-I PRS and Lipoprotein(a) (Lp(a)). Panel A. Prevalence of CAD per percentile of the PRS (green dots) and Lp(a) (red dots) distributions calculated in the testing population. The horizontal red line represents the value of CAD prevalence (2.8%) at which the Lp(a) distribution reaches a value of about 100 nmol/L (in correspondence of the 85^th^ percentile of its distribution). The same CAD prevalence value is reached by the PRS distribution at its 63^th^ percentile. **Panel B.** Scatter plot of the percentiles of the PRS distribution plotted against the percentiles of the Lp(a) distribution. Red continuous line: linear regression of the scatter plot.

In order to avoid reverse causation, the analysis involving plasma risk factors (measured at the UK Biobank first assessment) used only incident CAD. Incident CAD cases were identified by extracting for each individual of the testing sample the date of the first CAD event. The incident status was attributed to CAD events that occurred after the date of UK Biobank enrollment assessment.

Of note, LDL-cholesterol, total cholesterol and Apolipoprotein B levels for individuals reported to use cholesterol-lowering medications have been corrected by a correction factor of 1.56, 1.37, 1.46, respectively [40].

PRS shows the highest odds ratio per standard deviation (OR-SD) (1.69 (1.64-1.74)) respect to any single lipoproteins considered and the ratio between total cholesterol-HDL and ApoB-ApoA (Table 4).

**Table 4:**
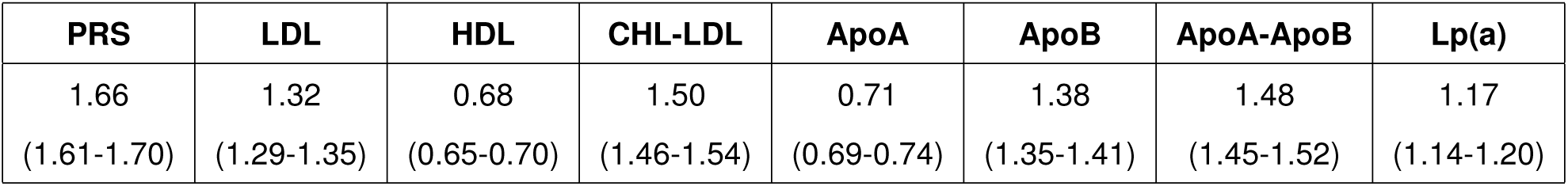
Odd Ratios per standard deviation for PRS and Plasma risk factors.

| PRS | LDL | HDL | CHL-LDL | ApoA | ApoB | ApoA-ApoB | Lp(a) |
| --- | --- | --- | --- | --- | --- | --- | --- |
| 1.66<br>(1.61-1.70) | 1.32<br>(1.29-1.35) | 0.68<br>(0.65-0.70) | 1.50<br>(1.46-1.54) | 0.71<br>(0.69-0.74) | 1.38<br>(1.35-1.41) | 1.48<br>(1.45-1.52) | 1.17<br>(1.14-1.20) |

#### 2.3.6 SCT-I PRS and age of CAD onset

Inouye et al [24] reported lower separation between PRS distributions of UK Biobank incident cases and controls compared with prevalent cases and controls. Since prevalent cases could have been characterised by a lower age of onset, they suggested a link between age of CAD onset and PRS score. We explore this hypothesis dividing the UK Biobank testing dataset in ten years age groups from 40 to 80 years (i.e. 40-50, 50-60, 60-70, 70-80) and calculating the AUC for each age group with a logistic regression including PRS, the 4 principal components of PCA ancestry, gender and genotyping array. A clear trend emerged from this analysis (Figure 10) where the higher AUC is observed in the younger group (40-50, AUC (CI):0.790 (0.774-0.809)) and steadily decreases with the increasing of age group down to the age group of 70-80 with AUC (CI): 0.691 (0.670-0.709). The genetic liability captured by PRS is therefore higher in early CAD cases, while late in life the manifestation of other risk factors could play a larger role than genetics. In light of this findings, PRS should be calculated in healthy individuals as early as possible to identify people who require stronger and earlier preventive interventions. PRS offers an invaluable opportunity to fight early CAD onset (i.e. <50 years), which is currently the most subtle and difficult to predict manifestation of the disease [41].

**Figure 10:**
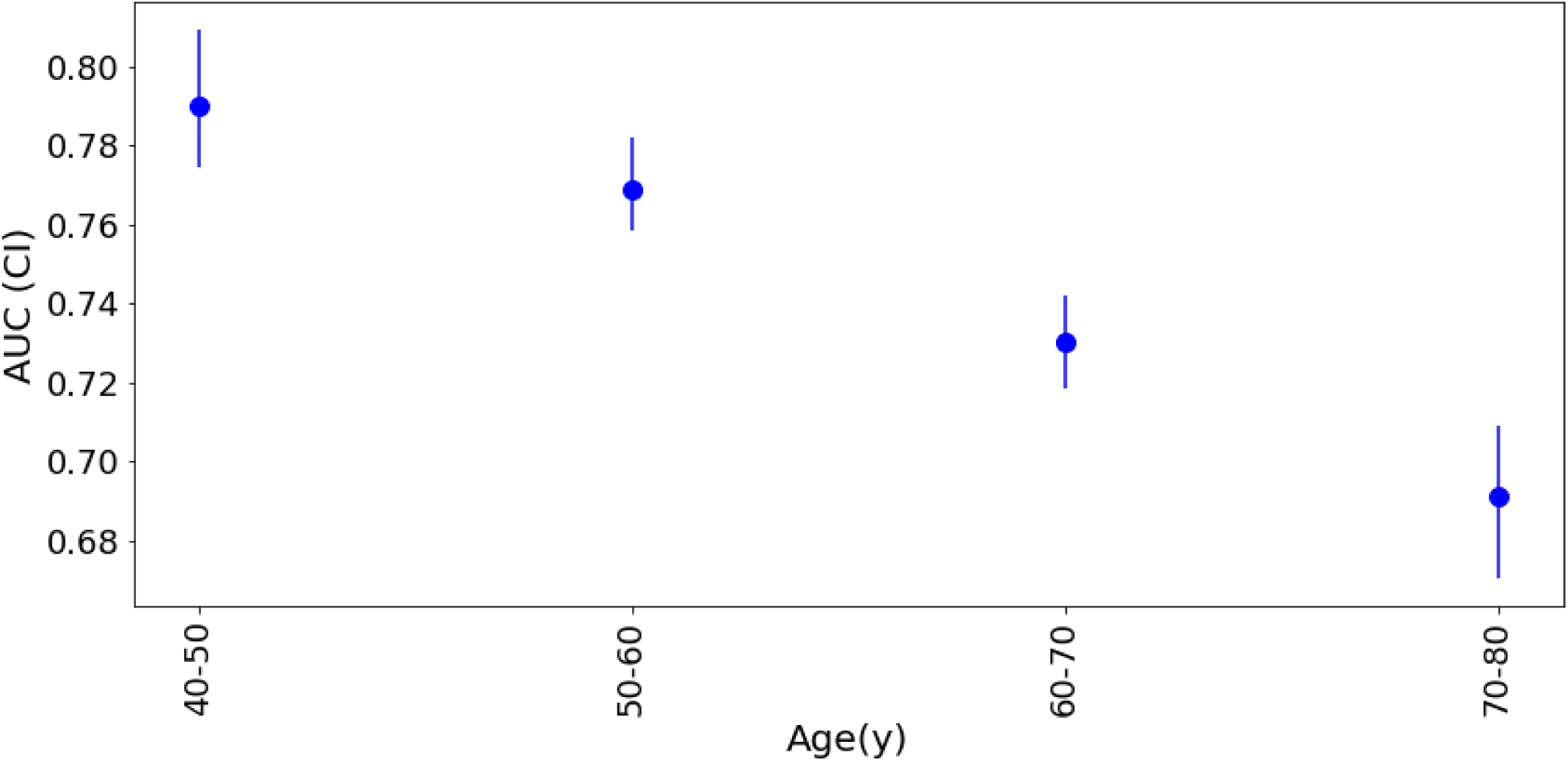
Predictive performance of the SCT-I PRS for different categories of age of CAD onset. Four testing datasets have been constructed using controls and CAD cases comprised between different age groups: 40 and 50 years; 50 and 60 years; 60 and 70 years; 70 and 80 years. The four datasets have been analyzed through logistic regression using the following covariates: SCT-I PRS, gender, genotyping array and the first 4 principal components (PCs) of ancestry. For each dataset, AUC and confidence intervals (IC) were determined

### 2.4 Breast Cancer

Breast cancer (BC) is the most common cancer diagnosed among women in Western countries [42]. The risk of developing BC is linked to both non-genetic and genetic factors. Non-genetic risk factors refer to not inherited nutritional, environmental (e.g., toxins), or pharmacological factors (e.g., hormone replacement therapy) [43–45]. From a genetic perspective, BC has a complex genetic architecture depending on two classes of genetic variations: rare mutations with high penetrance such as those of genes BRCA1 and BRCA2 [46, 47] and multiple common BC susceptibility loci that have been discovered through GWAS [15, 48, 49].

#### 2.4.1 Development of a BC PRS with improved predictive performance

We developed a BC PRS by applying the SCT algorithm described above and assessed its predictive performance in the testing dataset. The SCT PRS displayed higher predictive performance (AUC: 0.677, PPV at 3%: 20.20%) than other published BC PRS from Khera et al[22] (AUC: 0.65, PPV at 3%: 15.8%) and from Mavaddat et al[27] (AUC: 0.66, PPV at 3%: 18.0%) (Table 5). Notably, among the three PRS we compared, the SCT PRS used the largest number of SNPs (Table 5).

**Table 5:** List of the PRSs for BC assessed in this study. Khera refers to the PRS for BC developed by Khera et al^22^. Mavaddat refers to the PRS for BC developed by Mavaddat et al^27^. SCT refers to the PRS for BC developed with the SCT algorithm. For each PRS, Table 5 shows the number of genetic variants composing the PRS (**SNPs in PRS**), the predictive performances quantified as AUC values and 95% confidence intervals (**AUC (95% CI)**), the positive predictive values in the top 3% of PRS distributions (**PPV (3%)**), and the number of BC cases in the top 3% of PRS distributions (**Cases in top 3%**).

| PRS panel | SNPs in PRS | AUC (95% CI) | PPV (3%) | Cases in top 3% |
| --- | --- | --- | --- | --- |
| Khera | 5240 | 0.650 (0.640-0.658) | 15.80 | 721 |
| Mavaddat | 307 | 0.66 (0.651-0.670) | 18.0 | 819 |
| SCT | 577113 | 0.677 (0.667-0.686) | 20.20 | 921 |

#### 2.4.2 Predictive performance of the newly developed BC PRS

The following analyses are based on the SCT BC PRS that showed the highest predictive performance (Table 5). We computed the PRS of the individuals in the testing dataset and plotted the distributions of the scores for BC cases and controls (Figure 11A). The distributions are both gaussian, with BC cases showing a greater median PRS than controls (median: 0.51 and −0.04, respectively) and an AUC of 0.677.

**Figure 11:**
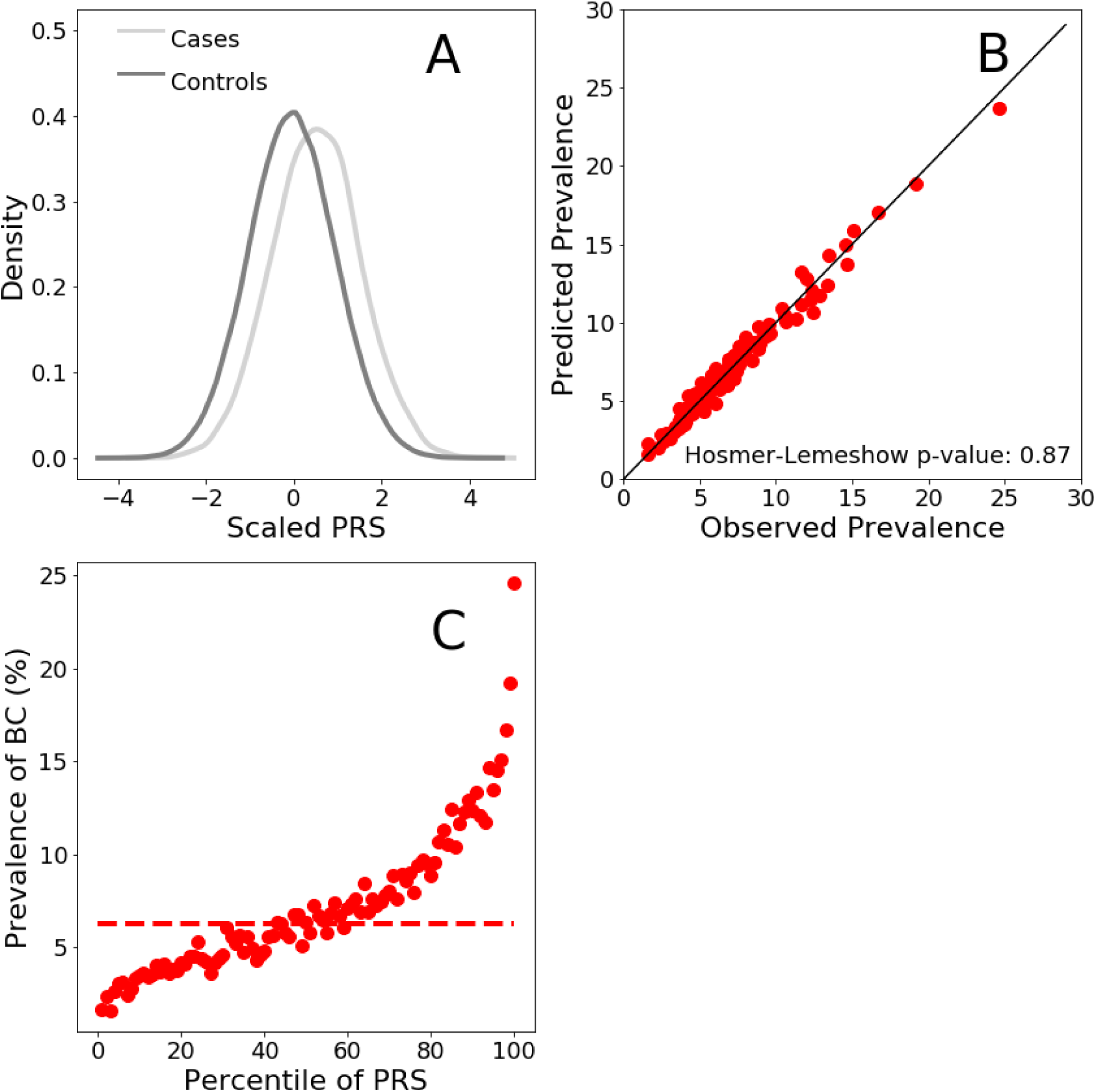
Risk for BC according to the SCT BC PRS panel. Panel A. Distributions (scaled to a mean of 0 and to a standard deviation of 1) of the PRS score for BC cases and controls in women testing dataset. **Panel B.** Comparison between the observed and predicted BC prevalence. Observed prevalence has been calculated as the prevalence of BC in the PRS distribution. Predicted BC prevalence was calculated for each individual using a logistic regression model with PRS as predictive variables. Within each percentile of the PRS distribution, BC probability was averaged and this returned the predicted prevalence of BC. **Panel C.** Dashed horizontal line: BC prevalence of the average of the PRS distribution (defined as between the 40% and the 60% percentiles)

We next assessed the ability of the SCT BC PRS to stratify BC risk for the female population in the testing dataset. As before, PRS distribution was divided into percentiles and we computed the prevalence of BC in each percentile. Disease prevalence is considered as empirical risk of developing BC. Risk stratification for women in the testing dataset is shown in Figure 11C. BC risk rises sharply as PRS percentile increases, ranging from 1.64% to 24.6%, for the lowest and highest percentiles respectively.

We estimated the relative increase in BC risk between the top 5% and the average (average of the percentiles comprises between 40% 60%, dashed line in Figure 11C) of PRS distribution. For the female testing population, the relative risk in the top 5% is 2.9 times higher than the average.

To assess the ability of the SCT BC PRS to predict BC risk in a testing population we compared the predicted with observed prevalence values. For all percentiles, predicted BC prevalence was plotted against the corresponding values of observed prevalence (Figure 11B). Figure 11B shows that the values of observed and predicted BC prevalence are in excellent agreement as demonstrated by the localization of the points of the bisector of the graph. The non-significant P value generated by the HL testing (Figure 11B) is a further confirmation of the good statistical agreement between predicted and observed prevalence values.

#### 2.4.3 PRS is more effective at predicting BC risk than family history

As for CAD, family history is a well-recognized risk factor of BC and prospective studies demonstrate a consistent association with the disease [50, 51]. Current prevention guidelines recommend the incorporation of family history into risk estimation models that guides treatment decisions for BC [52]. In order to evaluate the potential clinical utility of BC PRS, we assessed the risk stratification properties of BC PRS in a sub-population with family history of BC. Additionally, we compare the performance of PRS and family history in predicting the onset of BC.

We computed scaled distributions for BC cases and controls for those individuals in the testing dataset with at least one first-degree relative with a history of BC. Risk distributions for cases and controls are shown in Figure 12A. Both distributions are gaussian with cases having a higher median value than controls (median: 0.70 and 0.17, respectively). This suggests that the good discriminatory ability of the SCT BC PRS is maintained even in the population already considered at BC risk based on family history.

**Figure 12:**
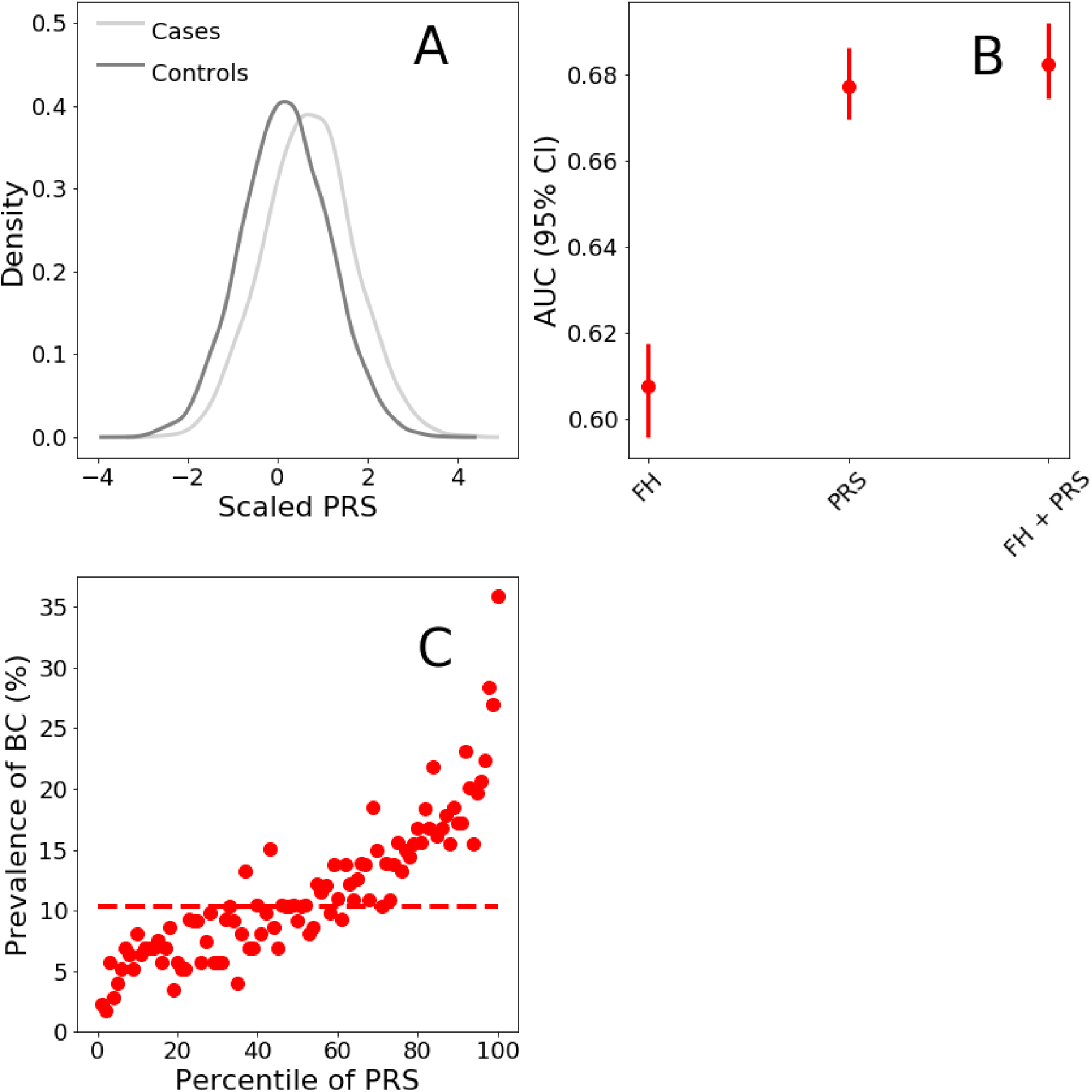
Risk for BC according to the SCT BC PRS panel in presence of Family history of BC. Panel A. Distributions (scaled to a mean of 0 and to a standard deviation of 1) of the PRS score for BC cases and control individuals with at least one first-degree relative with a history of BC. **Panel B.** AUC values were calculated on the testing dataset with logistic regressions using family history of BC (FH), PRS calculated with the SCT BC PRS (PRS), or both (FH + PRS) as explanatory variables. The response variable of the logistic regression model was absence/presence of BC. The logistic regression model comprised additional covariates as control variables such as: age, genotyping array, and the first 4 principal components (PCs) of ancestry. **Panel C.** Prevalence of BC per percentile of the PRS score distribution calculated in the testing population for women with at least one first-degree relative with history of BC. Dashed horizontal line: BC prevalence of the average of the PRS distribution (defined as between the 40% and the 60% percentiles).

We then evaluated the ability of the SCT BC PRS to stratify BC risk in the population with family history of BC. BC risk stratification for women with family history of BC are shown in Figure 12C. The SCT BC PRS is able to further stratify BC risk over a range of values comprised between 2.3% and 35.8%, for the lowest and highest percentiles, respectively. The observed prevalence is higher in women with family history than in the general population for any percentile considered. For women with family history of BC the relative risk in the top 5% is 2.6 folds higher than the average.

Lastly, we assessed the predictive performance of family history, BC PRS, and the combination of the two, by computing AUC. Figure 12B shows that the BC PRS displays a higher AUC value (AUC PRS 0.677 (0.667-0.686), AUC family history 0.606 (0.597-0.617) and when both risk factors are combined, the predictive performances improve (AUC 0.683 (0.673-0694).

### 2.5 Prostate Cancer

Prostate Cancer (PC) is the most common non-cutaneous cancer among men in the Western world [53]. Previous works estimated that more than 2000 common SNPs independently contribute to PC risk among populations of European ancestry [54]. While PRS have already been developed to predict PC risk, those PRS rely on small sets of GWAS-derived genome-wide significant SNPs [14, 55]. We therefore tested whether a large number of SNPs combined with the SCT algorithm could improve the predictive performance of previous PC PRSs.

#### 2.5.1 Development of a new PC PRS with improved predictive performance

We developed a new PC PRS by applying the SCT algorithm and assessed its predictive performance in the testing dataset. The new SCT PRS displayed higher predictive performance (AUC: 0.798, PPV at 3%: 19.8%) than the recently published PC PRS from Schumacher et al[22] (AUC: 0.774, PPV at 3%: 15.8%) (see Table 6). Notably, among the two PRS compared in Table 6, the SCT PRS was by far the one with the highest number of SNPs.

**Table 6:** List of the PRSs for PC assessed in this study. Schumacher refers to the PRS for PC developed by Schumacher et al^14^. SCT refers to the PRS for PC developed with the SCT algorithm. For each PRS, Table 6 shows the number of genetic variants composing the PRS (**SNPs in PRS**), the predictive performances quantified as AUC values and 95% confidence intervals (**AUC (95% CI)**), the positive predictive values in the top 3% of PRS distributions (**PPV (3%)**), and the number of BC cases in the top 3% of PRS distributions (**Cases in top 3%**).

| PRS panel | SNPs in PRS | AUC (95% CI) | PPV (3%) | Cases in top 3% |
| --- | --- | --- | --- | --- |
| Schumacher | 147 | 0.774 (0.766–0.782) | 15.8 | 598 |
| SCT | 682397 | 0.798 (0.787–0.807) | 19.8 | 752 |

#### 2.5.2 Predictive performance of the newly developed PC PRS

The following analyses are based on the SCT PC PRS that showed the highest predictive performance (Table 6). We computed the PRS of the individuals in the testing dataset and plotted the distributions of the scores for PC cases and controls (Figure 13A). The distributions are both gaussian, with PC cases showing a greater median PRS than controls (median: 0.70 and −0.04, respectively) and an AUC of 0.798.

**Figure 13:**
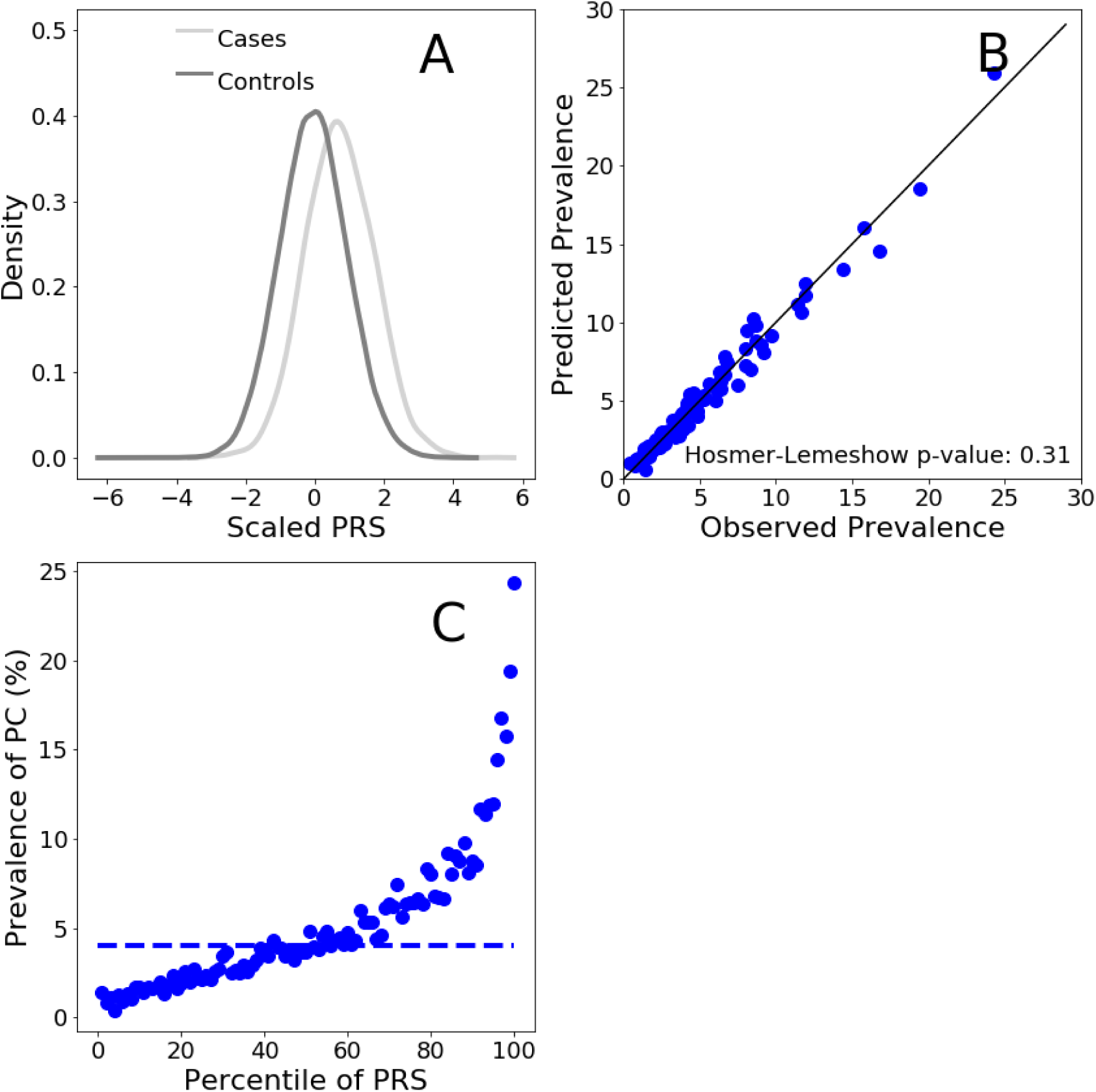
Risk for PC according to the SCT PC PRS panel. Panel A. Distributions (scaled to a mean of 0 and to a standard deviation of 1) of the PRS score for PC cases and controls in the male testing population. **Panel B.** Comparison between the observed and predicted PC prevalences. Observed prevalence has been calculated as the per-percentile prevalence of PC in the PRS distribution. Predicted PC prevalence was calculated for each individual using a logistic regression model with PRS as predictive variables. Within each percentile of the PRS distribution, PC probability was averaged and this returned the predicted prevalence of PC. **Panel C.** Prevalence of PC per percentile of the PRS distribution calculated in the men testing population. Dashed horizontal line: PC prevalence of the average of the PRS distribution (defined as between the 40% and the 60% percentiles)

We next evaluated the ability of the SCT PC PRS to stratify risk in men of the testing dataset. We divided the PRS distribution into percentiles and computed the prevalence of PC in each percentile. We used disease prevalence in the testing dataset as a measure of the empirical risk of developing PC. Risk stratification for men in the testing dataset is shown in Figure 13C. PRS stratify risk ranging from 1.43% to 24.3%, for the lowest and highest percentiles respectively, indicating a powerful clinical actionability for PC primary prevention.

We also estimated the relative increased risk, which is the ratio between the prevalence at the top 5% of the PRS distribution and the prevalence in the average of the distribution (defined as between the 40% and the 60% percentiles, dashed line in Figure 13C). For the men testing population, the relative risk in the top 5% is 4.1 times higher than the average.

To assess the calibration of the SCT PC PRS to predict PC risk in the testing population we compared the predicted with observed prevalence values. The values of observed and predicted PC prevalence are in excellent agreement as shown in Figure 13B. The non-significant P value generated by the HL test (Figure 13B) confirms that model is well calibrated.

#### 2.5.3 PRS is more effective at predicting PC risk than family history

As for CAD and BC, family history is a well-recognized risk factor of PC [56], we therefore considered the relationship between PRS and family history.

We computed scaled distributions for PC cases and controls for those individuals in the testing dataset with at least one first-degree relative with a history of PC. Risk distributions for cases and controls are shown in Figure 14A. Both distributions are gaussian with cases having a higher median value than controls (median: 0.90 and 0.15, respectively). This suggests that the discriminatory ability of the SCT PC PRS is maintained even in individuals already considered at PC risk based on family history.

**Figure 14:**
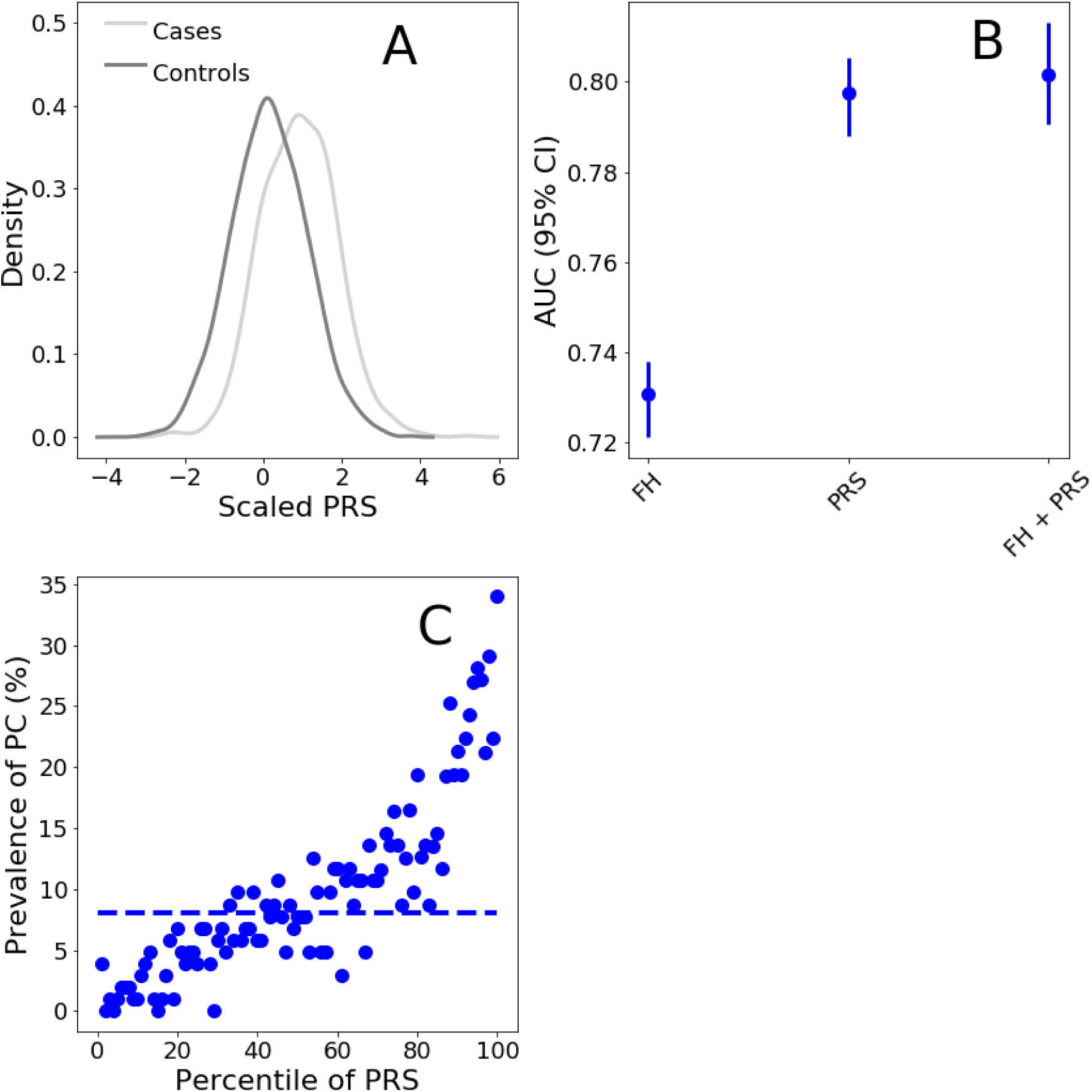
Risk for PC according to the SCT PC PRS panel in presence of Family history of PC. Panel A. Distributions (scaled to a mean of 0 and to a standard deviation of 1) of the PRS score for PC cases and control individuals with at least one first-degree relative with a history of PC. **Panel B.** AUC values were calculated on the testing dataset with logistic regressions using family history of PC (FH), PRS calculated with the SCT PC PRS panel (PRS), or both (FH + PRS) as explanatory variables. The response variable of the logistic regression model was absence/presence of PC. The logistic regression model comprised additional covariates as control variables such as: age, genotyping array, and the first 4 principal components (PCs) of ancestry. **Panel C.** Prevalence of PC per percentile of the PRS score distribution calculated in the testing population for men with at least one first-degree relative with history of PC. Dashed horizontal line: PC prevalence of the average of the PRS distribution (defined as between the 40% and the 60% percentiles).

We then evaluated the risk stratification of the SCT PC PRS in men with a family history of PC. PC risk stratification for men with family history of PC are shown in Figure 14C. Even with individuals considered at higher risk based on family history, the SCT PC PRS is able to stratify PC risk over a range of values comprised between 1.0% and 34.0%, for the lowest and highest percentiles, respectively. For men with family history of PC the relative risk in the top 5% is 3.1 folds higher than the average.

We computed AUC of family history, PC PRS, and the combination of the two. Figure 14B shows that the PC PRS displays a higher AUC value than family history (AUC PRS: 0.798 (0.787–0.807), AUC family history: 0.732 (0.722-0.743). Combining the two risk factors we obtain an AUC of 0.802 (0.793-0.813).

## 3 IMPLEMENTATION OF THE SAAS FOR GENOMIC RISK PREDICTION

### 3.1 SaaS Overview

To overcome current technological and practical challenges of implementing PRS in the clinic, we developed a Software as a Service (SaaS) for Polygenic Risk Score calculation. The SaaS for genomic risk prediction comprises a set of PRS panels from which an estimate of an individual’s risk of developing several polygenic diseases can be computed. Above, we demonstrated the predictive performance of 3 PRS panels for Coronary Artery Disease (CAD), Prostate Cancer (PC), and Breast Cancer (BC). In addition to those three, the SaaS utilised additional PRS Panels such as Atrial fibrillation (AF), Type 2 Diabetes (T2D), Type 1 Diabetes (T1D), Hypertension (Hyp), Inflammatory Bowel Disease (IBD) and Coeliac Disease (CD). Moreover new PRS panel can be added to the SaaS accepting as input a CSV file. In the following section we describe the SaaS pipeline.

### 3.2 The SaaS system: Genetic Data upload and conversion

The SaaS accepts genetic data in a variety of different formats. This expands the compatibility of the SaaS to any known microarray and Next-generation sequencing platforms. A genetic laboratory can upload the file containing the genetic information of an individual through a web interface with the secure FTPS protocol. Genetic data are accepted in the following formats:

- Standard Variant Call Format (VCF) files
- PLINK 1 binary (*.bed, *.bim, and *.fam)
- Oxford Format (*.gen, *.bgen, and *.sample)
- 23andme Text format files
- Non-standard custom text file formats

Data files are loaded to a shared Network file system and a new 16-cores Virtual Machine with 60 GB of RAM on a single tenant node generated for each analysis that needs to be performed. Data are analyzed on a single tenant node in order to guarantee efficient workload isolation and maximum security for the user. Once generated, each Virtual Machine activates a consecutive series of chained shell commands. In turn, each shell command triggers a series of either Python scripts or external software. The whole set of scripts representing the data conversion pipeline of the SaaS is listed below:

1. The input file provided by the user is read and converted to an internal format.
2. Genetic data are annotated with the appropriate dbSNP-ID, the latter belonging to either gene notation assembly GRCh37/hg19 or GRCh38.
3. Alleles of the samples provided by the SaaS user and those in the imputation reference panel must refer to the same strand for proper imputation (see below). However, the strand of the alleles varies depending on the genotyping and sequencing technology used. Therefore to identify the genetic variants in the samples that require a strand flip, the BEAGLE [57] strand check utility is used during this step.
4. For each genetic variant, the genotype is converted from a notation based on nucleic acid composition (A, T, C, G) to a binary ALT/REF notation.
5. Data from the 1000 Genome Project is used as reference panel to perform the conversion from A/T/C/G to ALT/REF notations.
6. Genetic data codified in the new ALT/REF notation are written in a series of files, one for each chromosome.

### 3.3 The SaaS system: Imputation

Genotype imputation is a process that allows an increase in the density of genetic data through the use of a statistical inferential procedure. In particular, imputation allows the estimation of uncalled genotypes and it is based on finding common haplotypes between an individual’s genome and a reference panel. Missing genotypes are then inferred from common haplotypes found in the reference set. This process leads to the estimation of the posterior probability distributions of the genotypes based on the available data. The SaaS utilised an imputation strategy that includes two reference panels: the 1000 Genome Project and the Haplotype Reference Consortium (HRC). The former has an higher number of SNPs (80M) but fewer haplotypes (2504) compared with the latter (SNPs 39M, haplotypes 64976). When there’s a match between SNPs in the two referenece panel the variants are imputed using the HRC to exploit the higher number of haplotypes available that can confer higher imputation accuracy. The software utilized in the imputation process is Beagle 5.0; it returns as outputs a series of vcf files, one for each chromosome. The final result of this imputation phase corresponds to the estimation of the genotype at 80 million genetic variants.

### 3.4 The SaaS system: Quality control of converted data

Converted imputed genomic files are subjected to a stringent quality control aimed at removing genetic variants of poor quality. The first quality control refers to the genotype probability of imputed genetic variants. Imputed genomic data display a triplet of values that refer to the probability of carrying combinations of the two alleles: two copies of the reference alleles, a copy of each allele, or two copies of the alternative allele. Genetic variants that display a maximal probability value below 0.8 are considered of poor imputation quality and thus are removed from the analyzed sample.

We performed a series of internal tests which showed that the predictive performance of a PRS remain stable up to a fraction of missing variants. For CAD, above a 10% missing variants threshold, the predictive power starts to deteriorate as the fraction of missing variants increases (Figure 15). Therefore in this step of the analysis, the number of missing genetic variants is estimated in each sample, and the sample is discarded if the fraction of missing variants is above the PRS specific missing variants threshold.

The complete series of quality controls is listed below:

1. Minor allele frequency: genetic variants with a low Minor Allele Frequency (MAF) are more prone to genotyping errors. For these reason, variants with MAF < 0.01 are removed from the sample.
2. Hardy - Weinberg equilibrium (HWE): genetic variants that display a deviance from HWE with a P value lower than 10*^−^*^6^ may indicate genotyping error and are therefore excluded from the analysis.
3. Ambiguous genetic variants: genetic variants with allele combinations A/T and C/G represent ambiguous variant because it is not possible to ascertain the strand of origin. For this reason ambiguous variants are removed from the analysis.
4. High Heterozygosity: Heterozygosity represents the proportion of heterozygous genotypes for a given individual. Deviations can indicate sample contamination. For this reason, Heterozygosity is computed for each individual under analysis and if this parameter deviates 3 Standard Deviation from the reference panel mean, the individual is removed from the analysis.

**Figure 15:**
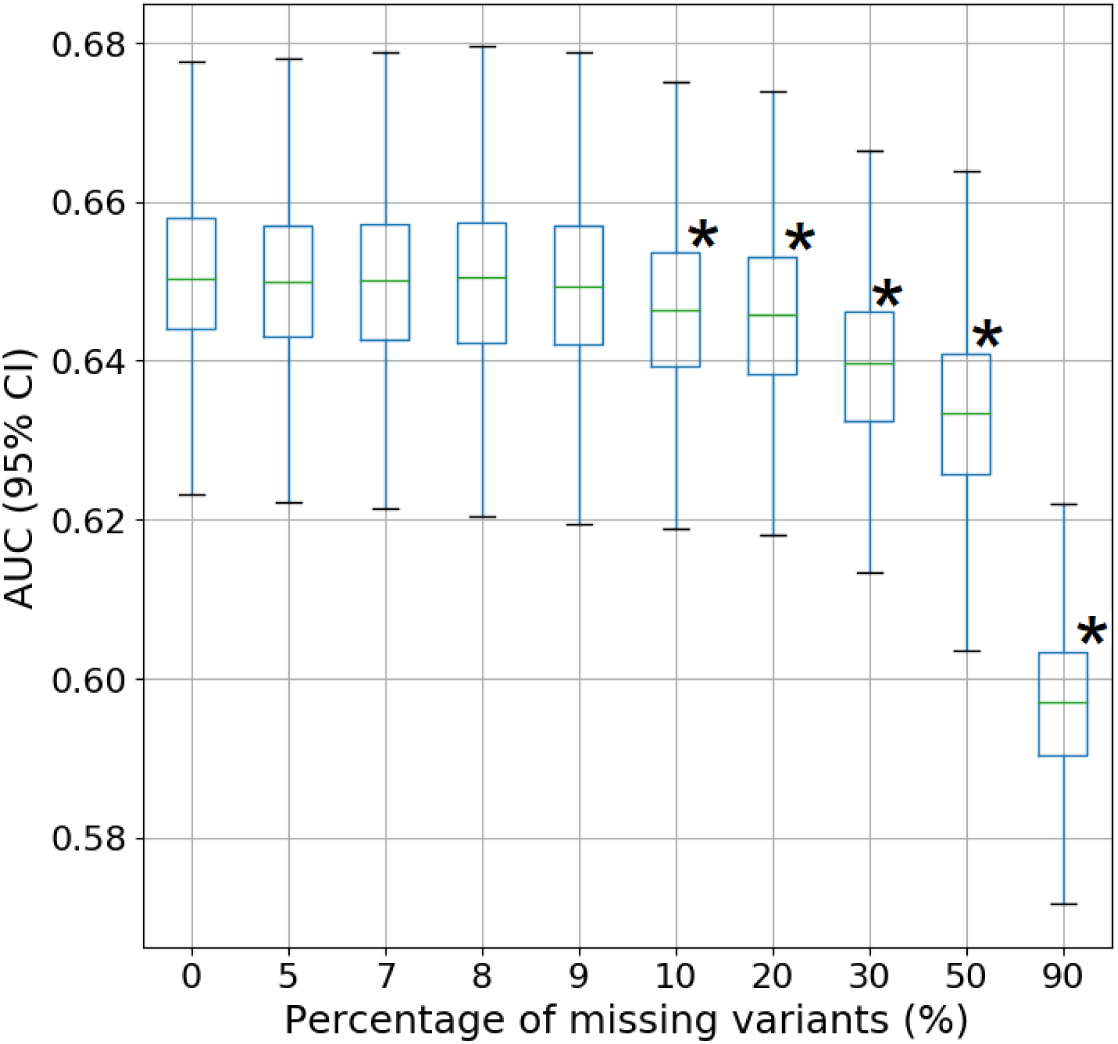
Dependence of the CAD PRS predictive performances on the fraction of missing genetic variants. A European population sample from UK Biobank consisting of 1470 CAD cases and 39000 controls was used for the analysis. The PRS panel for CAD from Khera et al^22^ (denoted as **Khera full** in Table 3) was used to calculate PRS values for each individual in the sample. PRS was used as the only predictive variable in a logistic regression model with binary CAD phenotype (0: control, 1:case) as response variable. The resulting predictive performances of the CAD PRS have been assessed by computing the AUC. The analysis has been repeated with different random fractions of missing genetic variants from the PRS panel. Asterisks denote the AUC distributions that display a statistically significant difference respect with the reference distribution (0% of missing variants), according to t-test analysis. Please note that the AUC value showed in this figure differs from the value reported in Table 2 and Table 3 because the latter has been calculated in a logistic regression with additional covariates such as age and gender.

### 3.5 The SaaS system: PRS calculation

The per-individual PRS is the final output of the analysis performed by the SaaS. For each disease considered in the SaaS, the corresponding PRS panel is characterized by three set of parameters: (1) A specific set of key SNPs; (2) A corresponding set of effect alleles that are statistically associated with the occurrence of each disease; (3) a corresponding set of effect sizes generated through the SCT algorithm. These PRS panels are used to calculate the per-individual PRS for each disease by summing the number of risk alleles in each genetic variant of the individual weighted by the corresponding value of the effect size. The predictive performances of the PRS for each disease are shown in section 3 for CAD, PC, and BC. The calculation of the PRS is based on a proprietary algorithm (written in Python 3) that utilizes as inputs the files generated by Beagle (Section 3.2) as well as a set of text files. For each individual and each disease, the corresponding value of the PRS score is calculated and compared with the average PRS score value of UK Biobank. This allows the estimation of a per-individual relative risk that is then used to identify individuals at high risk of developing the diseases. The final results of the SaaS are communicated, for each disease, in the form of a personalized report that contains information and plots that display the localization of the individual’s PRS compared to the reference population, the estimate relative risk of developing the disease as well as general guideline to be used for the correct interpretation of the results.

## 4 CONCLUSIONS

In this paper we described three PRS for complex diseases: CAD, BC and PC. We tested their prediction and risk stratification performances in the UK Biobank, which is the largest population dataset currently available. When compared with previously published PRS, the three PRS panels showed highest predictive performance(Tables 3, 5, and 6). Additionally, we demonstrated that PRSs for CAD, BC, and PC are able to identify a notable fraction of UK Biobank population (5%) with a 3 fold or higher increased risk of developing CAD, PC, and BC compared to the population average. The risk stratification ability of CAD, BC, and PC PRS is maintained even in individuals already considered at risk based on family history. Notably, CAD PRS has higher predictive power than lipoproteins routinely used as clinical risk factors. This implies that integrate PRS together with traditional risk factors in clinical risk models can enable physicians to more accurately quantify the risk of the diseases rendering more effective prevention and screening strategies. We developed a SaaS to perform PRS analysis, available to clinical laboratories and research groups as a fully automated, GDPR compliant and CE marked medical device. The SaaS calculates Polygenic Risk Scores for a large number of complex diseases and can analyse thousands of samples in parallel having the potential to improve health care prevention through its eventual large scale implementation into public health practice.

## Acknowledgements

We are grateful to all the volunteers that made the UK Biobank project possible and to the people involved in managing this extraordinary data resource, that was access as part of the Project 40692. We also thank Dr. Dina Radenkovich M.D. and Prof. Antonio Gaddi M.D. for their invaluable comments and inputs on the manuscript.

## Footnotes

* http://www.broadcvdi.org/informational/data

† https://www.cell.com/ajhg/fulltext/S0002-9297(18)30405-1

